# Data-Driven Equation-Free Dynamics Applied to Many-Protein Complexes: The Microtubule Tip Relaxation

**DOI:** 10.1101/2024.10.10.617682

**Authors:** Jiangbo Wu, Siva Dasetty, Daniel Beckett, Yihang Wang, Weizhi Xue, Tomasz Skóra, Tamara C. Bidone, Andrew L. Ferguson, Gregory A. Voth

## Abstract

Microtubules (MTs) constitute the largest components of the eukaryotic cytoskeleton and play crucial roles in various cellular processes, including mitosis and intracellular transport. The property allowing MTs to cater to such diverse roles is attributed to dynamic instability, which is coupled to the hydrolysis of GTP (guanosine-5’-triphosphate) to GDP (guanosine-5’-diphosphate) within the *β*-tubulin monomers. Understanding the equilibrium dynamics and the structural features of both GDP- and GTP-complexed MT tips, especially at an all-atom level, remains challenging for both experimental and computational methods because of their dynamic nature and the prohibitive computational demands of simulating large, many-protein systems. This study employs the “equation-free” multiscale computational method to accelerate the relaxation of all-atom simulations of MT tips toward their putative equilibrium conformation. Using large MT lattice systems (14 protofilaments × 8 heterodimers) comprising ∼21-38 million atoms, we applied this multiscale approach to leapfrog through time and nearly double the computational efficiency in realizing relaxed all-atom conformations of GDP- and GTP-complexed MT tips. Commencing from an initial 4 μs unbiased all-atom simulation, we interleave coarse projective “equation-free” jumps with short bursts of all-atom molecular dynamics simulation to realize an additional effective simulation time of 1.875 μs. Our 5.875 μs of effective simulation trajectories for each system expose the subtle yet essential differences in the structures of MT tips as a function of whether *β*-tubulin monomer is complexed with GDP or GTP, as well as the lateral interactions within the MT tip, offering a refined understanding of features underlying MT dynamic instability. The approach presents a robust and generalizable framework for future explorations of large biomolecular systems at atomic resolution.

**SIGNIFICANCE:** The dynamic instability of microtubules (MTs) is essential for a plethora of biological functions, from cell division to intracellular transport. Despite their importance, current computational models often struggle to handle the large-scale, long-term dynamics of all-atom simulations. This computational study employs the “equation-free” method to accelerate relaxation of the all-atom structures of MT tips in both GDP and GTP states which nearly halves the computational demands of simulating very large biomolecular systems. Our findings expose subtle but crucial structural differences between GDP-bound and GTP-bound MTs, particularly in the number of protofilament clusters, and is consistent with a relatively stronger lateral interactions in GTP-bound MT tips.

## 1. INTRODUCTION

All eukaryotic cells depend on microtubules (MTs) to perform a variety of essential cellular functions, including chromosome segregation during cell division, intracellular transport, and cell motility (1–4). Microtubules are cylindrical, hollow structures composed of *αβ*-tubulin heterodimer as their basic building blocks. As shown in **Fig. 1*a***, *αβ*-tubulin heterodimers are stacked longitudinally in a head-to-tail manner to form protofilaments (PFs), which are laterally associated to form the whole MT lattice, typically comprising 13 or 14 PFs (5–10). Both *α*- and *β*-tubulins of a free heterodimer can bind to guanosine-5’-triphosphate (GTP). The GTP molecule bound to *α*-tubulin at the interface of the *αβ*-tubulin heterodimer is nonhydrolyzable, whereas the GTP bound to *β*-tubulin is exposed to cytoplasm and can be hydrolyzed to guanosine-5’-diphosphate (GDP). The MT end with the GTP molecules of *β*-tubulins exposed to cytoplasm is referred to as the plus (+) end, where tubulin heterodimers exhibit a faster rate of polymerization and depolymerization. To fulfill their diverse biomolecular roles, MTs are highly dynamic, undergoing rapid and stochastic transitions from the polymerizing state to the depolymerizing state, known as the microtubule catastrophe (1,11,12). The inverse process called rescue refers to the switch back to the growing phase after the rapid shrinking phase from catastrophe. The collective process involving both catastrophe and rescue, known as the MT “dynamic instability,” is triggered by the hydrolysis of GTP bound to *β*-tubulin (1). It has been proposed that there is a “GTP cap” at the MT growing (+)-end, composed of an appreciable amount of GTP-bound tubulin heterodimers due to the lag between GTP hydrolysis and the recruitment of new heterodimers, which can protect MT from catastrophe (13–16). Upon the dissolution of the GTP cap, catastrophe begins and results in the release of mechanical strain in the MT lattice. This is considered to involve splaying of GDP-bound MT (+)-ends, resulting in a highly flexible ram’s horn-like structure (17–19). Recent studies also find the GTP-bound MT tip exhibits a splaying, ram’s horn-like structure (17–20). Accordingly, studying GTP and GDP bound MT systems can shed light on the equilibrium conformation and dynamics of MT during catastrophe and rescue, respectively. In addition to providing invaluable understanding of the MT dynamic instability, theoretical insights into the equilibrium structure and dynamics of GDP- and GTP-complexed MT tips can aid in the rational design of anti-cancer drugs targeting their malfunction (21–23).

**FIGURE 1.**
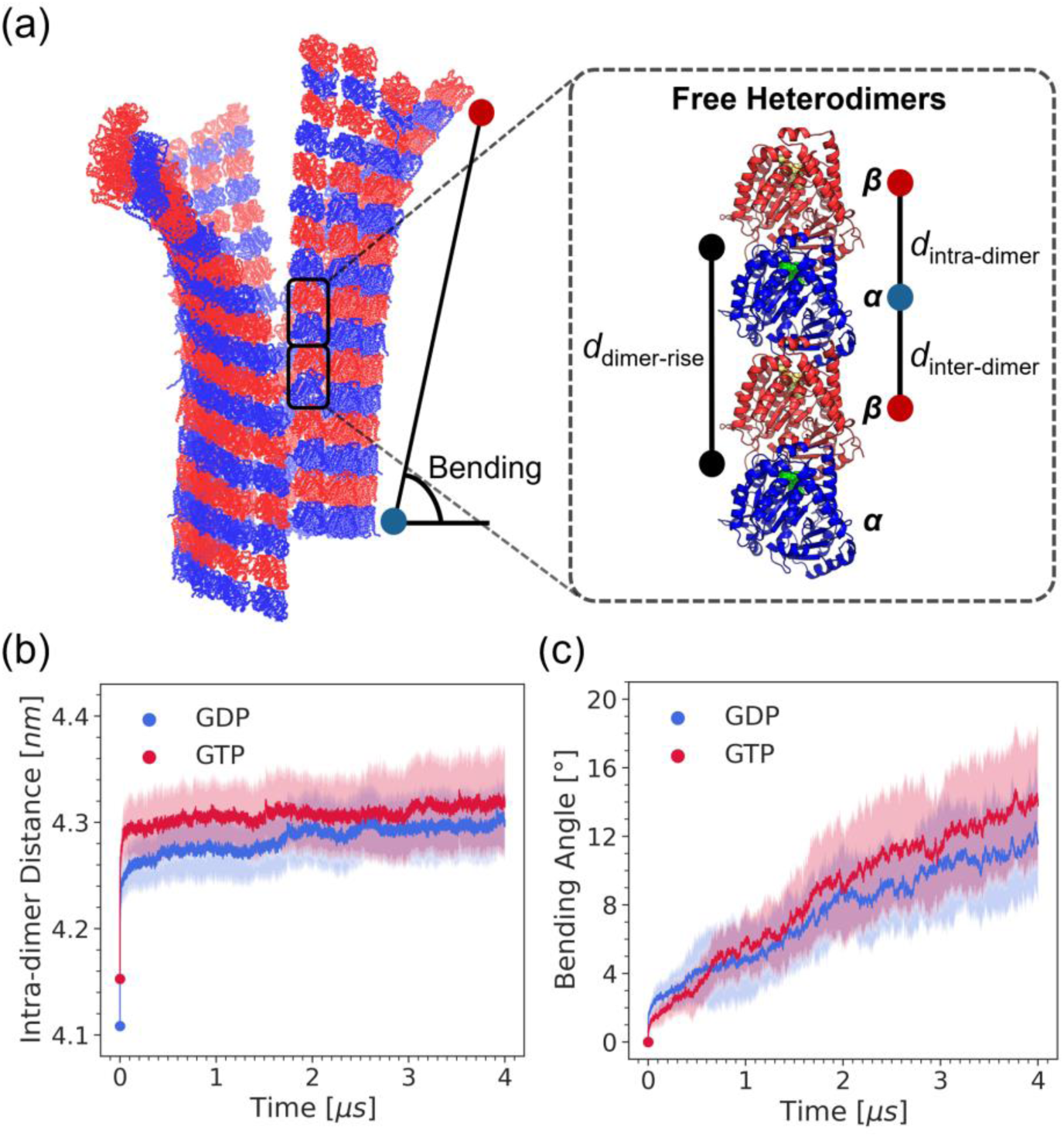
Slightly splayed MT tip structure simulated in this work illustrating the key parameters associated with the splaying process. (a) A 14 protofilaments (PF) × 8 heterodimers MT tip, with *α*-tubulin and *β*-tubulin backbone traces colored in *blue* and *red*, respectively. The dashed box shows two of the highlighted heterodimers with *α*-tubulin (*blue*) and *β*-tubulin (*red*) in cartoon representation. GDP (*green*) and GTP (*yellow*) nucleotides bound to *α*- and *β*-tubulin monomers, respectively are illustrated in van der Waals representation. The definitions of the vector or line segments used for calculating the four key structural parameters — bending angle, dimer rise (d_dimer-rise_), intra-(d_intra-dimer_) and interdimer (d_inter-dimer_) distances are also illustrated, where the solid circles at the end of vector or line segments represents the center of mass of either *α*-tubulin (*blue*) or *β*-tubulin (*red*). (b) Changes in the intra-dimer distances as a function of simulation time, with the GDP-bound MT colored *blue* and GTP-bound MT colored *red*. (c) Changes in the bending angles of GDP-bound MT (*blue*) and GTP-bound MT (*red*). In (b) and (c), the solid line indicates the average value over all the 14 PFs and the shaded region represents its standard deviation.

It is challenging to experimentally capture the detailed structure and mechanism of the splaying of MT tips because of its inherently dynamic nature. To this end, a series of theoretical and computational studies of these dynamic processes have sought to expose new understanding of the MT dynamic instability at a molecular level (19,20,24–31). Grafmüller and Voth (20) performed large-scale atomistic simulations for short PFs and showed the similar bending of the PFs in the GDP and GTP states. Tong and Voth (28) utilized all-atom (AA) molecular dynamics (MD) simulations to examine MT tip patches and lattices, highlighting the significant impact of the nucleotide states on MT dynamics. Gudimchuk et al. (26) used Brownian dynamics (BD) modeling to explore the mechanism of MT growth. They introduced an activation energy barrier to the lateral and longitudinal BD potentials, mimicking the electrostatic repulsion of surface negative charges and the entropic penalty for eliminating solvents. They found the activation barrier was a key parameter controlling MT force generation, which is critical for cellular processes like mitotic chromosome (7,32). Igaev and Grubmüller (27) employed long-time AA MD simulations on a single, two, or three laterally associated tubulin heterodimers in both GDP- and GTP-bound states to study their lateral interactions in the straight lattice conformation, using periodic boundary conditions to mimic the infinite protofilaments. Subsequently, Igaev and Grubmüller (29) expanded the system to a larger MT patch (14 PF ×6 heterodimers long) and performed extensive AA MD simulations to study the thermodynamics and kinetics of MT tips, demonstrating an energy barrier-governed mechanism of MT dynamic instability. While the computational studies applying AA MD simulations gained valuable insights into MT dynamic instability, the models employed often involved relatively small systems or were limited to short timescales due to the high computational demand. Accelerating the relaxation of MT tips using low-resolution coarse-grained (CG) methods or BD can circumvent this problem but often results in the loss of critical molecular-level details (33–36), including the subtle structures of the tubulin-tubulin interfaces, hydrogen-bonding information, the nucleotide binding pocket, and the heterogeneous contacts at the MT seam region. It remains a challenge to access the large-scale and long-time dynamics of the splaying process of MT tip at atomistic resolution near the equilibrium state.

In this work, our goal is to advance understanding of the differences in splaying of GDP- and GTP-complexed MT tip by using multiscale enhanced sampling techniques to nearly halve the computational cost of the expensive all-atom MD simulations and to access the putative relaxed equilibrium state in large-scale all-atom MD simulations. To this end, we used AA models of the MT tip with a lattice size of 14 protofilaments × 8 heterodimers long – comparable to that previously modeled by Igaev and Grubmüller (29) – and conducted AA MD simulations starting from a straight cylindrical conformation. With the application of approximately 56 million supercomputing CPU core hours on the Texas Advanced Computing Center (TACC) Frontera, we first performed 4 μs AA MD simulations for both GDP- and GTP-complexed MT tip lattice. Although we considered only one replica, it should be noted that the 4 μs of AA MD simulation represents an unprecedented length of trajectory for such a large system of MT tip based on our understanding (28,29), which contains approximately 21-38 million atoms. Several important parameters associated with MT tip splaying, including PF bending, dimer rise, intra- and interdimer distances, with a detailed definition in Zhang et al. (37), were calculated throughout the course of these initial 4 μs AA MD simulations. **Figures 1*b* and *c*** shows the time-series plots of PF bending and intra-dimer distances, which are commonly used metrics in cryo-electron microscopy (cryo-EM) experiments (37). **Supporting Information (SI) Figs. S1, S3 and S4** show the trends of all four parameters in these initial 4 μs long simulations. These plots show that the splaying process quantified by PF bending (**Fig. 1*c***) has not reached equilibrium for both the GDP- and GTP-complexed MT tips, indicating that the 4 μs long simulation is insufficient to access the equilibrium conformation of the splaying process. Additionally, the high flexibility and relatively small statistical differences between the GDP and GTP states of the MT tips make it challenging to discern their significant differences. Consequently, access to the equilibrium atomistic structures of both GDP- and GTP-complexed MT tip remained elusive despite these long 4 μs AA MD simulations.

To address the problem of the slow relaxation of the large, many-protein MT tip system towards its equilibrium state while retaining atomistic resolution, this work introduces an application of the “equation-free” multiscale simulation method (38–40). This approach uses coarse-projective integration through time within a subspace of collective variables (CVs), wherein short AA MD simulations sample the dynamical evolution of the system within that subspace of collective variables and inform a subsequent projective jump to leapfrog through time and thus realize significant computational savings. We have used the approach in this work here to achieve an effective simulation time of 5.875 μs of the dynamical evolution of GDP- and GTP-complexed MT tip at a ∼2-fold reduced computational cost relative to direct AA MD simulation. The large size of these ∼21-38 million atom simulations means that this saving equates to an approximate computational savings of ∼15 million CPU-hours. This technique has also enabled us to probe longer time scales than ever before in exploring the MT tip dynamics, and our results reveal the possible differences in the equilibrium structures of MT tips depending on whether *β*-tubulin monomer is complexed with GDP or GTP. Notably, we observed large differences in splaying of most of the PFs and their lateral interactions between the two nucleotide states. While the degree of splaying in the GTP-bound MT tip was higher, determining the statistical significance to compare with the experimental data require more than the single long simulation performed in this work. Nevertheless, the unprecedentedly long trajectories provide a refined understanding of the MT dynamic instability.

The remainder of this manuscript is organized as follows: first, we provide an overview of the “equation-free” multiscale technique applied to the large, many-protein MT-tip system. This is followed by the details of each of its operations — evolution of AA MD simulations (mature and drift), parameterization of the manifold by the selected CVs (restrict), learning and leap-frogging within the manifold using linear models (project), and backmapping from the parameterized manifold to AA space (lift) using steered-MD simulations (41). Then, we discuss the evolution and convergence of the CVs and show the structural features of the near equilibrium MTs in GDP and GTP states realized by the “equation-free” acceleration. Finally, we provide concluding remarks, highlighting the advantages and limitations of our approach in the acceleration of MT tip and similar large, many-protein systems using the “equation-free” multiscale method.

## 2. THEORY AND METHODS

### 2.1. Overview of the “equation-free” multiscale method applied to the MT tip system

The slow AA MD relaxation such as that of the intra-dimer distance and bending angle of MT tip (**Fig. 1*b*** and **Fig. 1*c***) suggests the presence of a slow subspace in which the dynamics of certain collective variables (CVs) evolve much slower than the remaining degrees of freedom in the system. Evolution of the slowest relaxing collective variables governs the approach to equilibrium, and accessing the equilibrium all-atom conformation using conventional molecular simulations can be computationally prohibitive especially for large, many-protein systems such as the MT tip containing ∼21-38 million atoms. The high computational cost can however be reduced by performing simulations at a coarse-grained resolution or by directly evolving the system in the slow subspace using the equation governing the slow observables in the MT tip system. With the coarse-graining approach, we lose access to important fine-grained information such as subtle structures of the tubulin-tubulin interfaces, hydrogen-bonding details, and the nucleotide binding pocket structure with coarse-grained simulations. On the other hand, determining the equations governing the slow observables in the MT tip system can be challenging. Kevrekidis and co-workers (38–40) pioneered the “equation-free” multiscale method that avoids the need for knowing the manifold subspace or the closed-form expressions governing the dynamics over it for accelerating the dynamics of a system while retaining access to the fine-grained information. Accordingly, we adapt this multiscale “equation-free” approach to address the challenge in accelerating the slow relaxation of the large, multi-million atom simulations of MT tip towards their equilibrium all-atom conformation as a function of whether the β-tubulin monomer is complexed with GDP or GTP.

Briefly, the “equation-free” approach refers to a generic data-driven multiscale modeling method that enables acceleration of microscopic dynamics by iteratively learning the emergent macroscopic dynamics from the microscopic system. **Fig. 2** illustrates the framework of the “equation-free” approach applied to accelerate the relaxation of MT tip dynamics towards their all-atom equilibrium conformation. The essence of this method is to conduct short all-atom MD simulations of the system to inform coarse-projective integration in time in a subspace of CVs. The new state point in CV space is then used to initialize and relax new all-atom MD simulations, and the process is repeated. By leapfrogging the all-atom simulation time in each coarse projective jump, this multi-scale approach can therefore realize large savings in computational effort and accelerate convergence towards equilibrium. The “equation-free” process can be formalized in five steps (**Fig. 2**): (1) lift, (2) mature, (3) drift, (4) restrict, and (5) project.

**FIGURE 2.**
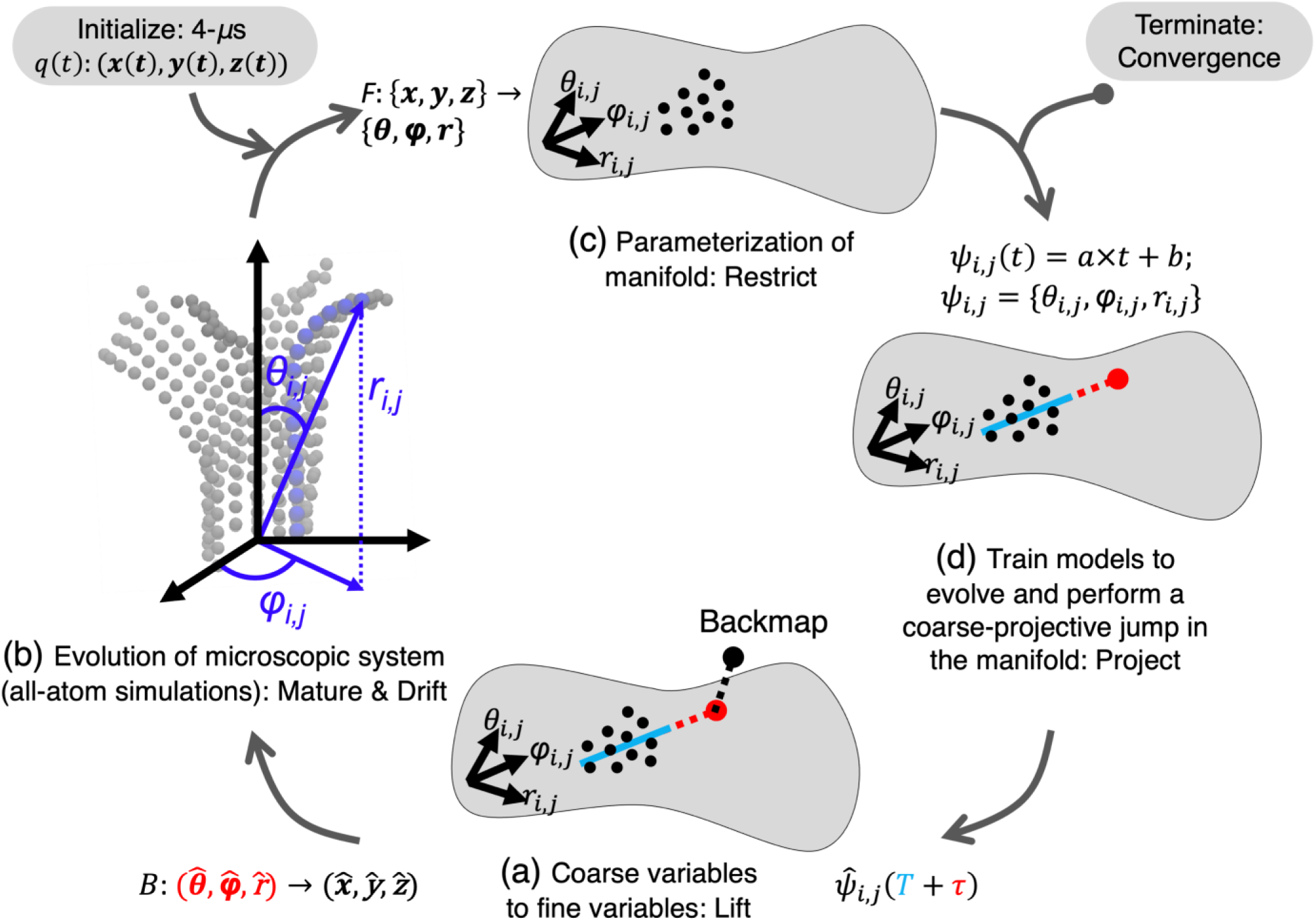
Illustration of the application of the “equation-free” multiscale method to accelerate the relaxation of GDP- and GTP-complexed MT tip systems. The method involves five sequential operations namely, (1) lift, (2) mature, (3) drift, (4) restrict, and (5) project. We started with the restriction step by mapping an initial 4 μs AA MD simulation trajectory into a CV subspace for both GDP- and GTP-complexed MT tip system. We selected the polar coordinates (*r_i,j_*, *θ_i,j_*, *φ_i,j_*) of the COM of each tubulin monomer *j* ∈ {1..16} in the MT tip as the CVs because they capture the bending (*θ_i,j_*) and twisting (*φ_i,j_*) angles as well as the lattice parameter (*r_i,j_*) changes in each PF *i* ∈ {*a*..*n*}. (a) A slightly splayed MT tip structure with only the COM of the tubulin monomers shown using gray spherical beads. The CVs ψ_*i*,*j*_ = (*r*_*i*,*j*_, θ_*i*,*j*_, φ_*i*,*j*_), are illustrated using the plus end β-tubulin monomer *j* of one of the PF *i* beads colored in blue. We define the polar coordinate system with the x- and y-coordinates of the origin at the center of the circle formed by the COM of the bottommost 14 *α*-tubulins. The z-coordinate of the origin corresponds to the minimum z-coordinate of the COM of the same bottommost 14 *α*-tubulins. We define the z-axis to be the principal axis of the straight, cylindrical MT lattice and the *x*-*y* plane to be the plane oriented perpendicular to the MT wall. Mature and drift steps refer to the evolution of the MT tip system at AA resolution using explicit-water MD simulations with AA resolved *α*- and *β*-tubulin monomers at each (*r_i,j_*, *θ_i,j_*, *φ_i,j_*) obtained from the lift operation. (b) After the AA MD simulations, the AA trajectories from the drift step are mapped into the CV subspace (*F*: {**x**, **y**, **z**} → {**r**, **θ**, **φ**}), known as the restriction operation. We use the trajectory length of *T* = 1 μs corresponding to the 3 to 4 μs simulation time of the initial AA MD simulation to initiate this restriction step, our starting point of the “equation-free” method. (c) The drift in the CV space from the restriction step informs the CV dynamics and the new state point in the CV space, which we learn (solid blue line) and predict (dashed red line) after time τ by training linear models ψ_*i*,*j*_(*t*) = *a* × *t* + *b* with parameters *a* and *b* on the drift trajectories of each CV ψ_*i*,*j*_(*t*): {*r*_*i*,*j*_(*t*), θ_*i*,*j*_(*t*), φ_*i*,*j*_(*t*)} of length *T*, obtained in the previous iteration. This refers to the projection step. (d) We then lift or backmap the new predicted state point in the CV space 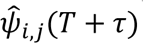 in the projection step after time τ to the all-atom space 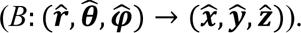 For this lifting process, we use steered MD simulations in implicit solvent. After this step, we place the system back under explicit solvation and start a new iteration by starting with the mature operation. We repeat the five operations until the convergence of the slow CVs, after which we terminate the process.

Lifting (**Fig. 2*d***) represents the initiation operation, where an initial or an evolved point in CV space is mapped to the all-atom space. This is a one-to-many mapping because many distributions in all-atom space could correspond to the same distribution in the CV space. Mature and drift operations (**Fig. 2*a***) refer to the relaxation and evolution of the system using all-atom MD simulations that are launched from the all-atom configurations obtained via the lifting operation. The resulting relaxed all-atom trajectories mapped into the CV space describes the restriction operation (**Fig. 2*b***). The coarse-projective jump to a new state point in the CV space based on the drift of the system in the CV space from the restriction operation is known as the projection operation (**Fig. 2*c***).

In the following, we first present the details of the 4 μs AA MD simulation that is used for initiating the “equation-free” multiscale approach for accelerating the relaxation of GDP- and GTP-complexed MT tip system towards their equilibrium conformational states. We then present the details of each of the five iterative “equation-free” operations by starting with the restriction step that includes selection of CVs for parameterizing the MT tip manifold. We end this section with the details of the convergence analysis.

#### 2.1.1. Initiation: All-atom MD simulation setup and details of the initial 4 μs simulations

Two cryo-EM crystal structures of MTs comprising 3 PF ×2 heterodimers in GDP-bound (PDB 6DPV) and guanylyl-(*α*-*β*)-methylene-diphosphonate (GMPCPP)-bound (PDB 6DPU) states were used to construct the initial all-atom structures for the MD simulations (37). Missing amino acids were modeled using MODELLER (42). To explicitly study the dynamics of the MT tip in the GTP-bound state, the nonhydrolyzable GTP analog GMPCPP was converted to GTP by substituting an oxygen atom for the CH_2_ linking group between the *α*- and *β*-phosphate.

To construct the larger 14 PF × 8 heterodimer MT tip (**Fig. 2*a***) lattice of both GDP and GTP states studied in this work, we leveraged the cylindrical symmetry of the original 3 PF ×2 patch structure from the cryo-EM structures using a procedure similar to that implemented by Tong and Voth (28). Full MTs were built by first extending longitudinally one heterodimer unit at a time using the alignment feature of VMD (43), and then extending laterally one PF at a time, to capture as much of the MT intrinsic curvature as possible. Specifically, the bottommost tubulin heterodimers of the 3 PF × 2 MT patch structure from the cryo-EM were aligned to the topmost tubulin heterodimers of another 3 PF ×2 MT patch, so that we got a new 3 PF ×3 MT patch. Similarly, the left PF of a MT patch was aligned to the right PF of another MT patch to extend a MT laterally. Finally, a seam region is formed between the 1^st^ and the 14^th^ PF, which has a longitudinal offset of three tubulin monomers. **SI Fig. S2** shows the resulting column structure of an MT tip used for both nucleotide states.

To allow for sufficient space of 2 nm along each axis for MT bending and avoid significant interactions between periodic images, we placed the cylindrical MT tip lattice structure in a simulation box of size 49.5 nm ×49.5 nm ×82.2/84.2 nm for MTs in GDP/GTP states, respectively with the z-axis of the box aligned along the cylindrical axis of the 14 PF ×8 MT tip lattice. We used TIP3P water model for solvating the system and neutralized it with 150 mM KCl. To accommodate the increase in MT tip splaying during the simulation, we periodically extended the simulation box three times by adding more water molecules and the corresponding amount of KCl to the system. The MTs were also rotated to position the splaying PF clusters diagonally within the cubic box to maximize space utilization. In total, the initial cylindrical structure of GDP and GTP simulation systems with water and ions contained 20,483,435 and 20,981,887 atoms, respectively. Towards the end of the “equation-free” process, the final near equilibrium structures of the MT tip system contained 35,189,277 and 38,452,523 atoms for GDP and GTP system, respectively. Detailed numbers of water molecules and ions in the initial and final systems are provided in **SI Table S1**.We employed the CHARMM36m force field with CMAP correction (44), with Particle Mesh Ewald for determining the long-range electrostatic interactions (45). Energy minimization was conducted in two stages using the gradient descent algorithm, with a maximum step size of 0.001 nm for the first stage and another 0.02 nm for the second stage. The system was pre-equilibrated at 1 atm and 310 K in the constant *NVT* ensemble for 100 ps, followed by equilibration in the constant *NPT* ensemble for 500 ps with the position restraints gradually removed from 1000 to 200 kJ/mol/nm^2^. To study the dynamic properties of the MT plus end and mimic the bulk environment within the MT lattice, position restraints were applied to all heavy atoms of the 14 *α*-tubulins at the bottom layer in the production *NPT* run (**SI Fig. S2**). Pressure and temperature were controlled by using the Parrinello–Rahman barostat (46) and canonical velocity-rescaling thermostat (47). The production runs of both GDP and GTP-complexed MT tip systems were performed for 4 μs using leap-frog integrator with a timestep of 2 fs. During the 4 μs production run, when the simulation box size was increased and water was added to the system to accommodate the increase in MT tip dimensions during splaying, we applied the same initial energy minimization and equilibrium protocol to the system before continuing the simulation. GROMACS version 2019.6 (48) was used for performing the energy minimization, equilibration and the production all-atom MD simulations. We used an average of 196 nodes each containing 56 Intel Xeon Platinum 8028 (“Cascade Lake”) cores on the TACC Frontera supercomputer for performing these all-atom MD simulations, which took ∼106 days for 4 μs for MT tip in each nucleotide state corresponding to a total parallel compute time of ∼28-million CPU-hours.

#### 2.1.2. Restrict: CV selection and mapping from all-atom space to CV subspace

The choice of CVs in the “equation-free” method for parameterizing the slow manifold remains a challenging problem similar to the selection of CVs in CV biasing enhanced sampling methods (49). In principle, appropriate CVs could be learned from the data within the so-called “equation-free / variable-free” paradigm (50), but prior structural and computational biology work have suggested a set of critical structural parameters responsible for the slow relaxation of the MT system (27–29,37). These include the bending (*θ_i,j_*) and twisting (*φ_i,j_*) angles of the PFs shown in **Fig. 2*a*** as well as the lattice distances (d_intra-dimer_ and d_inter-dimer_) illustrated in **Fig. 1*a*** and **SI Fig. S1**. For this reason, we selected (*r_i,j_*, *θ_i,j_*, *φ_i,j_*) coordinates of the center-of-mass (COM) of each tubulin *j* ∈ {1..16} in a PF *i* ∈ {*a*.. *n*} in the polar coordinate system as the CVs (**Fig. 2*a***) for parameterizing the MT tip manifold. These interpretable CVs are both easy to implement and capture the system’s principal motions. Specifically, *θ_i,j_* and *φ_i,j_* capture the bending and twisting motions; while *r_i,j_* helps in restraining the lattice parameters of the MT tip system. We defined the polar coordinate system with the x- and y-coordinates of the origin at the center of the circle formed by the COM of the bottommost 14 *α*-tubulins. The z-coordinate of the origin corresponds to the minimum z-coordinate of the COM of the same bottommost 14 *α*-tubulins. Because position restraints were applied to the bottommost 14 *α*-tubulins for mimicking bulk environment, the selected origin also remains largely fixed during the simulations. Further, we defined the z-axis to be the principal axis of the straight, cylindrical MT lattice and the *x*-*y* plane to be the plane oriented perpendicular to the MT wall (**Fig. 2*a***). Within this polar coordinate system, the trajectory of MT tip system can therefore be described on the manifold by the tuple (*r*_*i*,*j*_(*t*), θ_*i*,*j*_(*t*), φ_*i*,*j*_(*t*)), forming a 3*N*-dimentional vector of the COM of *N* = 14 × 16 = 224 tubulins at time *t*: {*r*_1_(*t*), θ_1_(*t*), φ_1_(*t*),…, *r*_*N*_(*t*), θ_*N*_(*t*), φ_*N*_(*t*)}, where the indices *i* and *j* are flattened for vectorization. In the tuple (*r*_*i*,*j*_(*t*), θ_*i*,*j*_(*t*), φ_*i*,*j*_(*t*)), *r*_*i*,*j*_(*t*) ∈ ℝ^+^, θ_*i*,*j*_(*t*) ∈ [0, π], and φ_*i*,*j*_(*t*) ∈ [0, 2π) with *r*_*i*,*j*_referring to the distance between the COM of tubulin *j* in PF *i* and the defined origin at simulation time *t*. θ_*i*,*j*_(*t*) and φ_*i*,*j*_(*t*) represent the bending and twisting angles of COM of tubulin *j* in PF *i* with respect to the straight MT wall, respectively.

Restriction (**Fig. 2*b***) refers to the mapping of the MT tip system from the all-atom space (**x**, **y**, **z**) to the CV subspace manifold (**r**, **θ**, **φ**) using the mapping function *F*. In particular, the mapping function *F* involves the evaluation of the 3*N*-dimensional vector {*r*_1_(*t*), θ_1_(*t*), φ_1_(*t*),…, *r*_*N*_(*t*), θ_*N*_(*t*), φ_*N*_(*t*)} from the all-atom MD simulations trajectory collected during drift operation of the previous iteration, with the starting drift data of *T* = 1 μs extracted from 3 to 4 μs of the initial 4 μs simulation conducted prior to the application of the “equation-free” method.

#### 2.1.3. Project: Coarse-projective integration in CV subspace

In the project stage of the “equation-free” method, the drift of the CVs ψ_*i*,*j*_(*t*): {*r*_*i*,*j*_(*t*), θ_*i*,*j*_(*t*), φ_*i*,*j*_(*t*)} in the restriction step informs the coarse-projective integration in time in the CV subspace (**Fig. 2*c***). The tracked CVs can be interpreted as the training dataset for learning the emerging CV dynamics of the MT tip system for extrapolating its evolution within the CV subspace. Projection therefore serves the crucial step in the “equation-free” method for realizing computational accelerations by “leapfrogging” to a predicted future state of the system without direct simulation of the intervening dynamics. In this work, we employed linear models ( ψ_*i*,*j*_(*t*) = *a* × *t* + *b*) for learning the dynamics of each of the CVs ψ_*i*,*j*_(*t*): {*r*_*i*,*j*_(*t*), θ_*i*,*j*_(*t*), φ_*i*,*j*_(*t*)}, as shown in **Fig. 2*c*** because of its simplicity and the relatively linear trends of CV trajectories during the drift operation at each iteration. We used the `curve_fit` tool of scipy library 1.11.2 (51) with Python 3.9.15 to train the linear model parameters (**a**, **b**) by using a least square minimization approach.

To determine the trajectory training window time *T* for learning the CV dynamics in each iteration and estimate the resulting error in predicting the CVs after a coarse-projective jump of time τ, we performed retrospective analysis using the initial 4 μs trajectory collected prior to the application of the “equation-free” method. In brief, this analysis involves training linear models over different trajectory slices of different lengths T within the initial 4 μs trajectory and comparing their predictions after various jump times τ with that observed in the initial 4 μs trajectory. From this retrospective analysis (**SI Fig. S19**), we found that the error in the linear model extrapolations 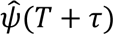 could be reduced to less than the standard deviations of the CVs observed in the 4 μs trajectory by selecting an appropriate training window time *T* of the drift trajectory and coarse-projective jump time τ.

In the first iteration, because we had access to the 4 μs trajectory, we trained the linear models over the trajectory window of time *T* = 1 μs corresponding to the 3 to 4 μs simulation time and performed a projective jump of τ = 0.5 μs. From the retrospective analysis, the mean error over all *i* and *j* between the predictions from the linear model 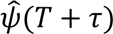 and the observed CV values ψ(*T* + τ) for a training window size of *T* = 1 μs and a coarse-projective jump of τ = 0.5 μs are (0.68±0.31)°, (1.51±0.92)°, and (0.15±0.12) nm for the *r*, θ, and φ CVs, respectively for the GDP MT tip system. For the same *T* = 1 μs and = 0.5 μs, the mean errors for the GTP MT tip system are (0.80±0.36)°, (1.79±0.99)°, and (0.19±0.09) nm for the *r*, θ, and φ CVs, respectively. These errors lie within the standard deviation of the CVs averaged over all *i* and *j* observed in the initial 4 μs trajectory (**SI Fig. S19**). For the subsequent “equation-free” iterations, the stride of each jump was reduced to τ = 0.125 μs, a conservative choice made to keep the extrapolation error low (**SI Fig. S19**). This was because of the smaller trajectory training window size of *T* = 0.25 μs collected during the drift step of the subsequent “equation-free” method iterations.

In principle, we note that the “equation-free” method allows for deviations from the CV subspace manifold during the coarse-projective jump and therefore, the extrapolations need not be accurate because upon an appropriate maturation time and with multiple configurations from the lift operation, the system can heal from the error during the projection step and relax on the CV free energy surface. However, because of the high computational cost associated with the large, many-protein MT tip system, and gradual maturation time, running multiple-replicas during lift operation is not so feasible and therefore necessitates a conservative coarse-projective jump time τ.

#### 2.1.4. Lift: Mapping from CV subspace to all-atom space

After making a jump in the CV subspace, a corresponding adjustment in the space of fine variables – that is the all-atom coordinates of MTs – should be applied. **Fig. 2*d*** illustrates the lift step, which can be conceived of as the inverse operation to restriction, where the predicted CVs 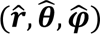 in the projection step are transformed from the CV subspace manifold to the all-atom space 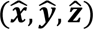 using the backmapping function *B*. In this work, we only backmap the positions of the MT tip system without solvent or salt ions. We note that even the cylindrical structure used for launching the initial 4 μs simulation prior to the application of “equation-free” method can be perceived as a lift operation from the CV subspace with ψ_*i*,*j*_: {*r*_*i*,*j*_, θ_*i*,*j*_, φ_*i*,*j*_} corresponding to that of the all-atom structure built from the cryo-EM 3 PF ×2 patch. The backmapping function *B* is a one-to-many function because many distributions in all-atom space could correspond to the same distribution in the CV space. In this work, steered-MD simulations were applied to drive the transformation from CV subspace 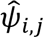 to the all-atom space 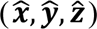 after the projection operation (41). The steered-MD simulation involves gradual application of a harmonic potential to guide an initial all-atom MT structure with its CV subspace location 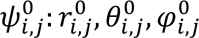 towards the projected configuration in the CV subspace with 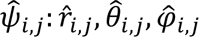 over a simulation time *t*. The harmonic bias potential can be expressed as 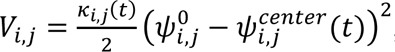, where κ (*t*) and 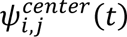 refer to the force constant and the target CV reference center of the harmonic potential at simulation time *t*, respectively. We scaled κ_*i*,*j*_(*t*) and 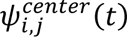 linearly over the simulation time *t* from 0 to 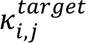 and 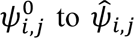, respectively. The two hyperparameters 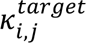 and simulation time *t* were adjusted for gradually transforming the initial all-atom MT structure with CV subspace location 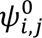 to a target location 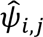 while also ensuring that the stability of the critical lattice parameters of MT d_intra-dimer_ and d_inter-dimer_ were within the bounds by comparing their distribution against the initial 4 μs trajectory obtained prior to the application of the “equation-free” method.

For both GDP and GTP-complexed MT, we found a 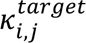 of 50,000 kJ/mol/nm^2^ over a simulation time *t* of 200 ps to be appropriate for all three CVs ψ_*i*,*j*_: {*r*_*i*,*j*_, θ_*i*,*j*_, φ_*i*,*j*_} in most of the “equation-free” iterations. In some iterations, we found a more gradual steering rate to 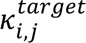 of 50,000 kJ/mol/nm^2^ over a simulation time 300-400 ps to be better based on the stability of the critical lattice parameters d_intra-dimer_ and d_inter-dimer_ of MT tip. In each of the “equation-free” iterations, we used the last all-atom configuration from the drift step as the initial all-atom MT structure with CV subspace location 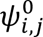 for starting the steered-MD simulations.

We implemented the CHARMM36m force field with a generalized Born implicit solvent model representation with the dielectric constants of solute and solvent set to 1 and 78.5, respectively to speed up these lifting calculations using steered-MD simulations. A Nosé-Hoover thermostat (52,53) implemented in OpenMM v8.0 (54) with leapfrog integrator using a timestep of 2 fs was applied to maintain the temperature at 310 K using a coupling frequency of 0.1 ps^-1^. We patched OpenMM with the PLUMED v2.7.3 (55,56) enhanced sampling library to apply the harmonic bias forces during the simulation. In each iteration and for both GDP and GTP-complexed MT tip system, we performed the steered-MD simulation for an additional 200 ps after κ_*i*,*j*_(*t*) reached 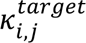. These simulations of 400 ps containing ∼1.5M atoms took ∼18 hours on 1 node with 1 V100 GPU and 40 Intel(R) Xeon(R) Gold 6148 CPUs.

#### 2.1.5. Mature and drift: Healing and evolution using all-atom MD simulations

Following the lifting stage, we proceed with the mature operation (**Fig. 2*a***), where the lifted all-atom MT tip structures **q**: (**x**, **y**, **z**) undergo a relaxation for allowing the system to “heal” from any deviations from the free energy surface of the manifold subspace during the projection operation. This mature operation is also crucial for adding back in the explicit solvent and ions omitted during the lift operation and in alleviating distortions introduced during the steered-MD. The input structures for this stage were obtained from the last frame of the steered-MD simulation in the previous stage. Due to changes in the MT tip dimensions during projection operation, a new simulation box was defined with a length suitable for the potential conformation change of the MT tip. Solvation and neutralization procedures were identical to the initiation step. Similarly, two stages of energy minimization and equilibration in the constant *NVT* and *NPT* ensembles were performed using the same scheme as in the initiation stage for this mature step.

At the end of mature step, we performed 0.25 μs of production run for evolving the system, which we refer to as the drift operation (**Fig. 2*a***). We set this drift simulation time of *T* = 0.25 μs to judiciously use the computational resources and keep the simulation wall time to less than about 9 wall-clock days. Subsequently, the 0.25 μs trajectory was passed to the next restriction operation and the remaining “equation-free” operations were repeated until convergence of the CVs. Details of the all-atom MD simulations during the mature and drift operations are identical to that of the initiation operation.

#### 2.1.6. Termination of the “equation-free” method: Convergence of the bending and twisting motions

In total, we performed four rounds of the equation-free approach, which involved a collective AA MD simulation time of 1 μs and a net coarse-projective jump of 0.875 μs for both GDP and GTP-complexed MT tip systems, corresponding to an approximately 2-fold computational acceleration and an estimated computational savings of ∼15 million CPU-hours. The termination criteria was based on the convergence of the collective time averaged *r_i,j_*-*θ_i,j_* and time averaged *r_i,_*_j_-φ_i*,j*_ values for a given PF *i*. Specifically, we monitored the convergence of bending motion of each PF by calculating the Fréchet distance between the curves formed by the 16 time averaged (*r_i,j_*,*θ_i,j_*) coordinates corresponding to the 16 tubulins *j* for each of the 14 PFs *i* in an iteration of the “equation-free” method and that observed in the initial 4 μs all-atom trajectory. Similarly, we calculated the Fréchet distance between the curves formed by the 16 time averaged (*r_i,j_*, φ_i*,j*_) coordinates in an iteration of the “equation-free” method and that observed in the initial 4 μs all-atom trajectory for measuring the convergence of the lateral twisting motion of each of the 14 PFs. The Fréchet distance estimates the similarity of two curves retaining consideration of the ordering of the coordinates constituting the curves (57–59), and serves as a suitable measure for estimating the convergence of the bending and twisting motions of the PFs in MT tip with the iterations in the “equation-free” method. In brief, the Fréchet distance captures the minimum path between two curves by traversing both the curves in the same direction without backtracking. Colloquially, it can be explained as the minimum leash length required for a person and a dog to walk along their respective paths without backtracking until their end points. We employed the discrete variant of the Fréchet distance (57) that can be determined using the recursive relation, 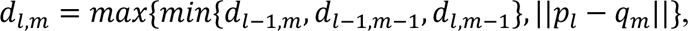, with *l* and *m* incremented from low to high values. Here, ||*p*_*l*_ − *q*_*m*_|| denotes the Euclidean distance between the points *p*_*l*_and *q*_*m*_of the two curves *P* and *Q,* respectively, with *P* formed by *N* {*p*_1_…*p*_*N*_} coordinates and *Q* formed by *M* {*q*_1_…*q*_1_} coordinates.

We projected the 16 time averaged polar coordinates *r_i,j_*-*θ_i,j_* and *r_i,_*_j_-φ_i*,j*_ into Cartesian space for each PF *i* prior to the calculation of the Fréchet distance between the curve formed in each iteration and that observed in the initial 4 μs all-atom trajectory. The Python library `frechetdist` 0.6 (60) was employed for computing all the Fréchet distances.

## 3. RESULTS AND DISCUSSION

### 3.1. An approximately 2-fold acceleration in MT tip relaxation dynamics using the “equation-free” multiscale method

Our goal in this work is to develop a method to accelerate the relaxation of GDP and GTP-complexed MT tip system at an all-atom resolution towards their equilibrium state using molecular simulations for gaining insights into the microtubule “dynamic instability” phenomena. We show that the “equation-free” multiscale based technique can aid in realizing this goal by boosting the computational speed by ∼2-fold, which for the large, many-protein MT tip system with ∼21-38 million atoms equate to a savings of 8.5 million CPU hours for every 1 *μ*s reduction in all-atom simulation time for the MT tip system in each nucleotide state. Briefly, the acceleration in the “equation-free” method is a result of leapfrogging over the all-atom simulation time with their corresponding coarse-projective jumps in a subspace of CVs as the guide. In this work, we elected to use (*r_i,j_*, *θ_i,j_*, *φ_i,j_*) coordinates (**Fig. 2*a***) of the COM of each tubulin *j* ∈ {1..16} in a PF *i* ∈ {*a*.. *n*} in the polar coordinate system as the CVs, which capture the important bending and twisting motions as well as correlate with the lattice parameters of the MT tip system.

We used the 4 μs all-atom unbiased simulation trajectories to seed the first restriction-projection-lift-mature-drift round of the “equation-free” method (**Fig. 2**) implemented using the selected CVs for GDP- and GTP-complexed MT tip systems shown in **Fig. 3**. In total, we performed four “equation-free” iterations with the termination criteria based on the convergence of the bending and twisting motions of the PFs in the MT tip. Specifically, we employed Fréchet distance (**Sec.** 2.1.6) between a bending or twisting curve observed in an iteration and that observed in the initial 4 μs all-atom trajectory to assess convergence. We considered 14 bending Fréchet distances corresponding to the bending motion of the 14 PFs *i* is formed by the 16 time-averaged (*r_i,j_*,*θ_i,j_*) coordinates corresponding to the 16 tubulins *j*. Similarly, we considered 14 twisting Fréchet distances associated with the 14 PFs *i* is formed by the 16 time-averaged (*r_i,j_*, φ_i*,j*_) coordinates corresponding to the 16 tubulins *j*.

**FIGURE 3.**
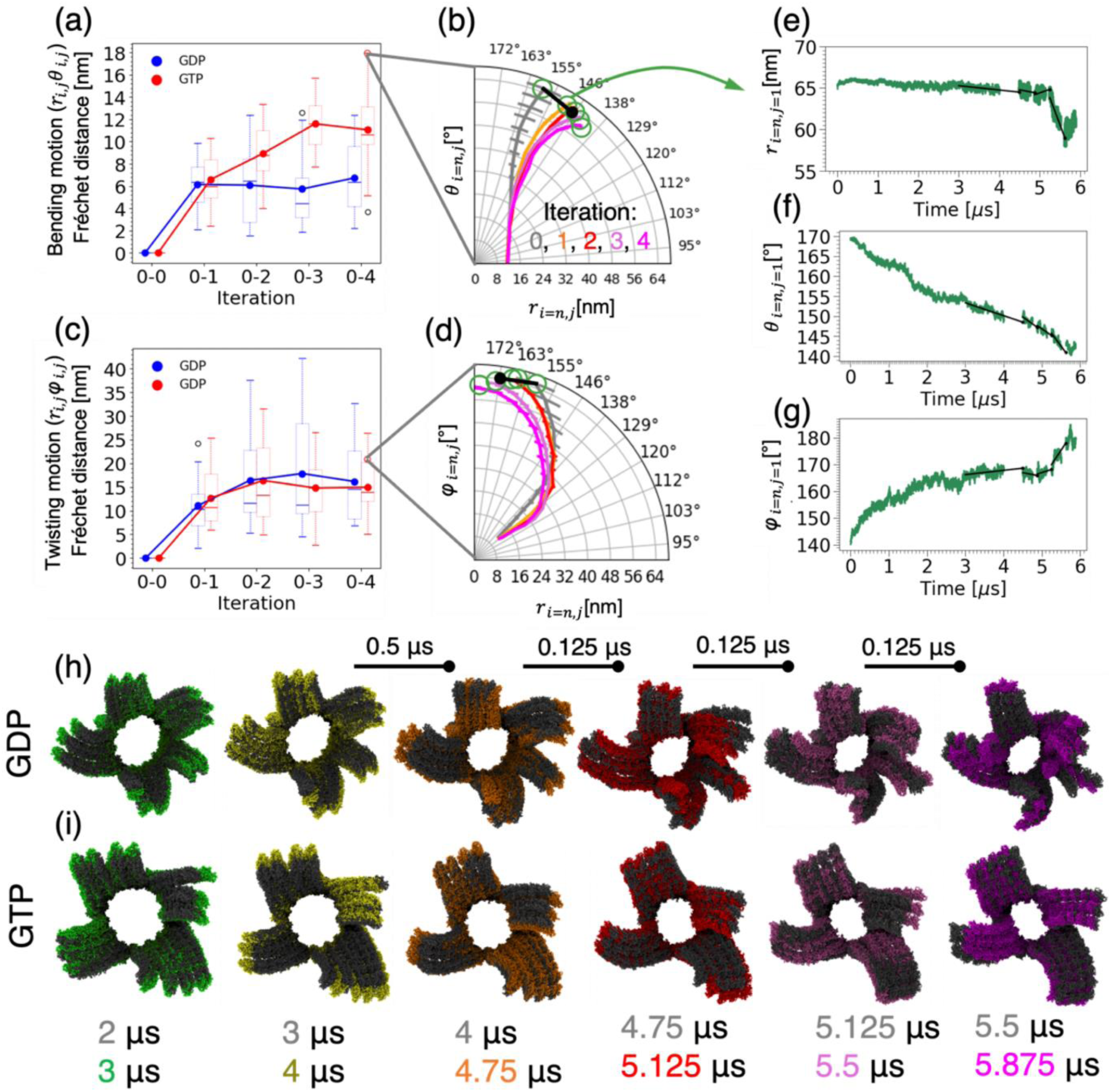
Accelerated convergence of the relaxation dynamics of GDP- and GTP-complexed MT tip with the “equation-free” method. We measured the convergence of the bending and twisting relaxation dynamics with “equation-free” iterations by monitoring the Fréchet distance between the curves formed by the respective 16 time averaged (*r_i,j_*,*θ_i,j_*) coordinates and the 16 time averaged (*r_i,_*_j_,φ_i*,j*_) coordinates for every given PF *i* ∈ {*a*.. *n*} in an iteration of the “equation-free” method and that observed in the initial 4 μs all-atom trajectory. (r*_i,j_*,θ*_i,j_*, φ*_i,j_*) are the CVs selected to parameterize the manifold, which represent the position of the center of mass of each tubulin monomer *j* in a PF *i* relative to a fixed reference point. (a) Changes in the Fréchet distance in nm between the 16 time-averaged r*_i,j_*-θ*_i,j_* coordinates in each of the 14 PFs in each of the iteration with respect to their corresponding coordinates in the initial 4 μs trajectory. Box plots at each iteration reveal the distribution of the 14 Fréchet distances that correspond to the 14 PFs. Solid lines are drawn by connecting the average of the 14 Fréchet distances at each iteration. Fréchet distance profiles of GDP- and GTP-complexed MT tip are colored in blue and red, respectively. (b) Illustrative curves formed by the 16 time-averaged r*_i=n,j_*-θ*_i=n,j_* coordinates for the PF *i*=n in GTP-complexed MT tip. Curves in the initial 4 μs are colored in gray and the four “equation-free” iterations are colored in orange, red, purple, and magenta, respectively. Error bars represent the standard deviation of the r*_i=n,j_*-θ*_i=n,j_* coordinates during the simulation. (c) Similar to (a) but shows the statistics from the 14 Fréchet distances obtained from the twisting curves formed by the 16 time-averaged r*_i,j_*-φ*_i,j_* coordinates. These Fréchet distances capture the relaxation of the lateral twisting of the 14 PFs. (d) Similar to (b) but corresponds to an example curve formed by the 16 time-averaged r*_i=n,j_*-φ*_i=n,j_* coordinates for the PF *i*=n in GTP-complexed MT tip. (e-g) Illustration of the application of the “equation-free” method along the CVs (*r_i=n,j=1_*, *θ_i=n,j=1_*, *φ_i=14,j=1_*) corresponding to the topmost *β*-tubulin *j*=1 of the plus end of the PF *i*=n. Linear models (solid black lines) are trained to learn the dynamical evolution of the CVs and perform an extrapolative coarse-projective jump (black circle markers) in the CV space. Snapshots illustrating the changes in the MT tip structure from plus (+) end perspective during the last 2 μs of the initial 4 μs trajectory and after applying the equation-free method for the (h) GDP- and (i) GTP-complexed MT tip. We illustrate the subtle but important structural changes during the initial 4 μs and at the end of the “equation-free” iterations relative a reference structure (gray). For the 3 μs (*green*) and 4 μs (*yellow*) structures from the initial simulation, the reference structures are 2 μs and 3 μs, respectively. The structures at the end of each of the four “equation-free” iterations are shown relative to the terminal structure of the previous “equation-free” iteration in orange, red, purple, and magenta, respectively.

In, **Fig. 3*a***, the distribution over the 14 bending Fréchet distances corresponding to the 14 PFs obtained from the bending coordinates (*r_i,j_*,*θ_i,j_*) are shown as box plots at each iteration for both GDP (blue) and GTP (red) complexed MT tip systems with their corresponding coordinates from the initial 4 μs data as the reference. Individual bending Fréchet distances for each of the14 PFs are shown in **SI Fig. S5**a and **SI Fig. S5**b. The mean of these 14 Fréchet distances (solid lines) reach a plateau after the first and third iteration for the GDP-(blue) and GTP-(red) complexed MT tip systems, respectively. This indicates that the bending motion of the PFs on average converged within the four iterations for both GDP- and GTP-complexed MT tip systems. An example change in the bending motion corresponding to the largest observed Fréchet distance, which is associated with a PF *i*=n (**SI Fig. S2**) of the GTP complexed MT system is shown in **Fig. 3*b***, where the 16 time averaged (*r_i=n,j_*,*θ_i=n,j_*) positions along with their standard deviations are plotted for both during the initial 4 *μ*s and the four “equation-free” iterations. **SI Fig. S6** and **SI Fig. S7** show the profiles for all the 14 PFs in both the GDP- and GTP-complexed MT tip systems. These profiles reveal a more detailed picture on the relaxation of the bending of each PF, where we observed 11 of the 14 PFs in GDP and all the PFs in GTP displayed a higher outward splaying at the end of four iterations relative to the initial 4 *μ*s trajectory with the “equation-free” iterations.

In addition to the slow relaxation of the bending of the PFs or *θ_i,j_* CVs, the lateral splaying or twisting of each PF or φ_i*,j*_ CVs is also a slow process. In **Fig. 3*c***, the distribution over the 14 twisting Fréchet distances corresponding to the 14 PFs obtained from the twisting associated (*r_i,j_*, φ*_i,j_*) coordinates are shown as box plots at each iteration for both GDP-(blue) and GTP-(red) complexed MT tip systems with their corresponding coordinates from the initial 4 μs data as the reference. Individual twisting Fréchet distances for each of the 14 PFs are shown in **SI Fig. S5**c and **SI Fig. S5**d. Here, the mean of the 14 Fréchet distances (solid lines in **Fig. 3*c***) reaches a plateau after 2 iterations for both GDP-(blue) and GTP-(red) complexed MT tip systems. This indicates that the lateral twisting motion of the PFs on average also converged within the four iterations for both GDP- and GTP-complexed MT tip systems. An example twisting profile change corresponding to the same PF *i*=n in the GTP complexed MT system highlighted in **Fig. 3*b*** for bending profile change with iterations is plotted in **Fig. 3*d***. **SI Fig. S8** show the lateral twisting profiles for all the PFs in both GDP and GTP complexed MT tip system in all the iterations. Here, the lateral grouping of neighboring PFs observed in the initial 4 μs all-atom trajectory are largely preserved during the equation-free iterations but the final iteration had a higher lateral twisting in the PFs for both GDP- and GTP-complexed MT tip systems.

Given the emergence of a plateau in the average of the 14 Fréchet distances of bending (solid lines in **Fig. 3*a***) and twisting profiles (solid line in **Fig. 3*c***) corresponding to the 14 PFs, we have terminated the “equation-free” method after four iterations and considered both the GDP- and GTP-complexed MT tip systems to have reached relaxed conformational states. Together with the four equation-free iterations and the initial 4 μs simulations, we have reached an unprecedent effective simulation time of 5.875 μs for both GDP and GTP complexed MT tip system. This implies the equation-free method resulted in a net savings of 0.875 μs since the net all-MD simulation time during the four iterations is 1 μs.

In **Fig. 3*e*-*g***, we illustrate the “equation-free” acceleration process that resulted in this 0.875 *μ*s savings by showing the restriction and projection step along (*r_i=n,j=1_*, *θ_i=n,j=1_*, *φ_i=n,j=1_*) corresponding to the top most *β*-tubulin *j*=1 of the plus (+)-end for the bending and twisting profile of the same PF *i*=n in GTP MT tip highlighted in **Fig. 3*b*** and **Fig. 3*d***, respectively. Detailed profiles for other values of *i* and *j* are shown in **SI Fig. S9**-**S14**. The four coarse-projective jumps in the CV subspace correspond to jump lengths of τ = 0.5 *μ*s, 0.125 *μ*s, 0.125 *μ*s, and 0.125 *μ*s, respectively with a total of 0.875 *μ*s. These jump lengths are determined based on the error estimates in our retrospective analysis (**Sec.** 2.1.3) of the CV predictions at different jump lengths τ with different training trajectory time lengths *T* for the linear models used for learning the CV dynamics.

The acceleration from the “equation-free” method and changes in the GDP- and GTP-complexed MT tip structure at each iteration can also be analyzed visually. In **Fig. 3*h*,*i***, we show the top views of the 3D MT tip structures at different effective simulation times. Changes in the structures from a side view at different effective simulation times and with other reference structures are shown in **SI Fig. S15**-**S18**. While the GDP- and GTP-bound MT tips display different PF bending and twisting profiles, these snapshots reveal a more detailed effect of the coarse-projective jumps in CV space (e.g., **Fig. 3*e*-*g***) directly on the all-atom space and the subsequent drift during the all-atom MD simulations. Consistent with the quantitative bending and twisting profiles (e.g., **Fig. 3*b*,*d***), we observe both outward and inward bending of the PFs and grouping of neighboring PFs in these detailed snapshots of the MT tip systems.

### 3.2. Differences in the near-equilibrium GDP- and GTP-bound MT tip conformations

If we consider the spread in the box plots of **Fig. 3*a*** and **Fig. 3*c***, we note that while the averages of the CVs tracking the bending and twisting motions of the PFs have converged, there are outliers which appear not to have fully relaxed within the four “equation-free” iterations. Furthermore, we have only considered one single trajectory for both GDP- and GTP-complexed MT systems, and therefore it is more challenging to assess the statistical significance of these specific trajectories. While we would like to continue additional rounds of “equation-free” acceleration and conduct independent replicas of these calculations, we are limited by the high computational cost of these large simulations. The “equation-free” approach enabled us to report a very long 5.875 μs long trajectory that has approached a relaxed conformational state for both the GDP- and GTP-complexed MT tip systems. We now investigate the structural differences between MTs in GDP- and GTP-bound states but restrict to the gross features because of our limited statistics. Specifically, we consider the root mean square deviation (RMSD) over the C-alpha (C*α*) atoms to analyze the MT tips at four levels: the entire microtubule, individual protofilaments, individual heterodimers, and individual tubulin monomers.

**Fig. 4*a*** shows the RMSD of the whole MT tip as a function of simulation time, using the straight cylindrical MT lattice as the reference structure for each state. The RMSD values for the GTP state (*red*) increase more rapidly and reach higher values compared to the GDP state (*blue*) during the initial 4 μs simulation and throughout the four stages of the “equation-free” extrapolation. The MT in the GDP-bound state shows a more gradual increase in RMSD. In both the cases, the RMSD curve reaches a plateau, but the higher RMSD of GTP-complexed MT tip relative to its initial straight MT lattice suggests that it is more dynamically flexible than the GDP-complexed MT tip, at least comparing the single replicas conducted for each system. However, these differences may not be statistically significant as noticed by Igaev and Grubmüller in their five independent runs of ∼2.4 μs of 14 PF × 6 heterodimers long MT tips in the GDP- and GTP-bound states (29).

**FIGURE 4.**
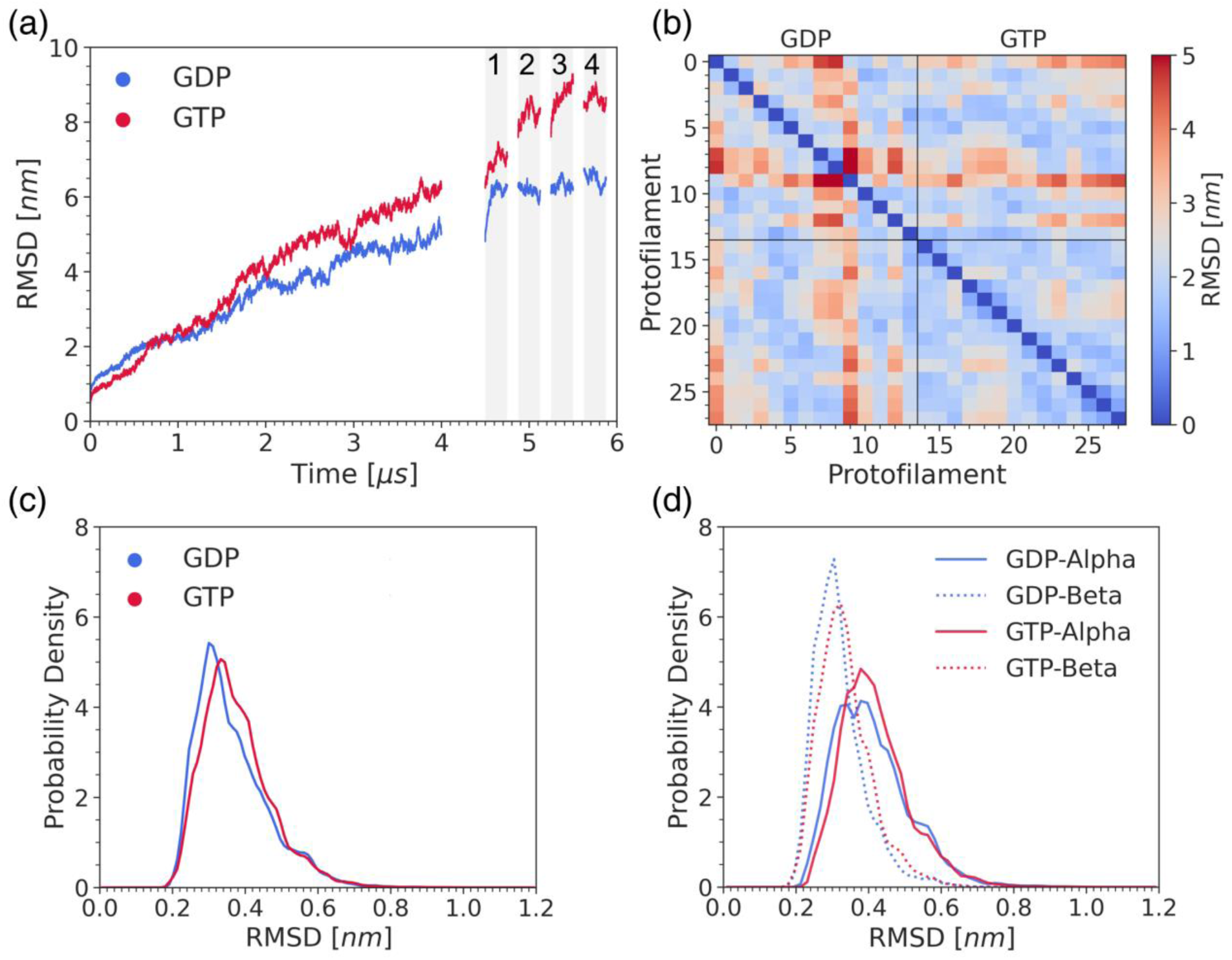
Structural comparisons between GDP- and GTP-complexed MT tip at the whole microtubule (MT), protofilament (PF), heterodimer, and tubulin levels. (a) MT level: RMSD curves of MTs in both GDP- and GTP-bound states varying with simulation time, using the initial straight MT structures as the reference. The *gray* shading denotes the four equation-free iterations. (b) PF level: Pairwise RMSD matrix between all 28 PFs of MTs in GDP and GTP states throughout the four equation-free trajectories. Ticks from 0 to 13 represent the 14 PFs of GDP-bound MT, while 14 to 27 indicate the 14 PFs of GTP-bound MT in ascending order. (c) Heterodimer level: distribution of the RMSD of tubulin heterodimers, with reference to the initial structure throughout the equation-free trajectories. (d) Tubulin level: distribution of the RMSD of tubulin monomers, with reference to the initial structure throughout the equation-free trajectories. In (a), (c), and (d), the GDP- and GTP-complexed RMSD profiles are colored in *blue* and *red*, respectively.

**Figure 4*b*** presents a pairwise RMSD heatmap between all 2 × 14 PFs of MTs corresponding to both GDP and GTP states. For this calculation, we excluded the initial 4 μs trajectory as part of the equilibration process and report the RMSDs between two PFs by averaging over the conformations obtained in the four equation-free iterations. The upper left block corresponds to the average RMSD of the PF-PF of GDP-complexed MT, and the lower right block represents RMSD between PF-PF of the GTP-complexed MT. Generally, the MT in GTP state exhibits lower RMSD values across each pair of PFs, indicating a larger similarity in the PF structures potentially because of a concerted motion of the PFs. In contrast, the MT in the GDP state displays a mixture of low-RMSD and high-RMSD pairwise distances, suggesting that the PFs in GDP-complexed MT split into a higher number of clusters with similar PF structure in each cluster. These observations are consistent with the quantitative lateral twisting angle profiles (**SI Fig. S8**) and the visual analysis in **Fig. 3*h*** and **Fig. 3*i***.

The distribution of RMSD values at tubulin level is shown in **Fig. 4*c***. Here, the RMSD of each heterodimer with reference to the heterodimer structure in the initial straight cylindrical MT lattice is calculated, except for the tubulin heterodimers at the bottom layer which are subjected to position restraints throughout the simulation. The MT in GTP state exhibits a marginally broader distribution than the GDP state, suggesting greater variability in the heterodimer structures along the MT tip. Further, the MT in the GDP-bound state has a statistically smaller (*p*-value = 0.037 < 0.05) average RMSD, reflecting more stable and consistent structural behavior in the heterodimer structure. The statistical differences between the averages are estimated using a one-tailed T-test by slicing the trajectory into five slices.

Lastly, the RMSDs at the individual *α*- and *β*-tubulin monomer level are shown in **Fig. 4*d***. As illustrated, the *α*-tubulin monomer has a larger RMSD values than *β*-tubulin for MTs in both GDP- and GTP-bound states. This indicates that *α*-tubulins display larger changes in its structure than the *β*-tubulin in both GDP- and GTP-complexed MT tip systems.

### 3.3. Differences in the effective interactions of the near-equilibrium GDP- and GTP-bound MT tip conformations

Lateral interactions are demonstrated to play key roles in regulating MT dynamic instability (12,61,62). For this reason, we also studied the differences in the lateral interactions of the near-equilibrium GDP- and GTP-complexed MT tip conformations by using the 0.25 μs trajectory of the final “equation-free” method iteration, which is the iteration closest to the structurally relaxed state. We caution, however, against over-interpretation of this analysis due to the fact that these results are based on a single trajectory replica for each system.

For studying the lateral interactions, we employed a heterogeneous elastic network model (HeteroENM) to parameterize the MT tips and estimate the strength of the effective interaction pairs, using the 0.25 μs trajectory from the last equation-free iteration. An interaction pair is considered effective if its spring constant is larger than 0.05 *k*_B_*T*/Å^2^ (63,64). For constructing the HeteroENM, each tubulin is coarse-grained into a single CG bead and positioned at the COM of its C*α* atoms. We analyzed all effective HeteroENM pairs, distinguishing between the seam and non-seam regions, as illustrated in **Fig. 5*a* and *b***. In **Fig. 5*a***, MT in the GTP-bound state exhibits fewer, yet, on average, stronger effective pairs with spring constants (0.64 ±0.29) kcal/mol·Å^2^ compared to that in the GDP-bound state with spring constants (0.31 ±0.22) kcal/mol·Å^2^ in the seam region (*p*-value = 0.049 < 0.05 for the one-tailed *T*-test). The MT in the GTP state also shows relatively sparse effective pairs, which aligns with the more pronounced splaying of the seam region in GTP-bound MTs in our simulations (**Fig. 3*h* and *i***, and **SI Fig. S17**). **Figure 5*b*** compares the effective interaction pairs of MTs in both GDP and GTP states in the non-seam regions, with average spring strengths of 0.33 kcal/mol·Å^2^ for the GDP state and 0.48 kcal/mol·Å^2^ for the GTP state (*p*-value = 0.007 < 0.01 for the one-tailed *T*-test). The results from two panels (**Fig. 5*a*** and **Fig. 5*b***) confirm that the GTP state MT tip shows stronger lateral interaction than the GDP state MT tip, and highlight the seam region as particularly vulnerable region to bending, which reinforces the arguments of the previous work by Tong and Voth (28).

**FIGURE 5.**
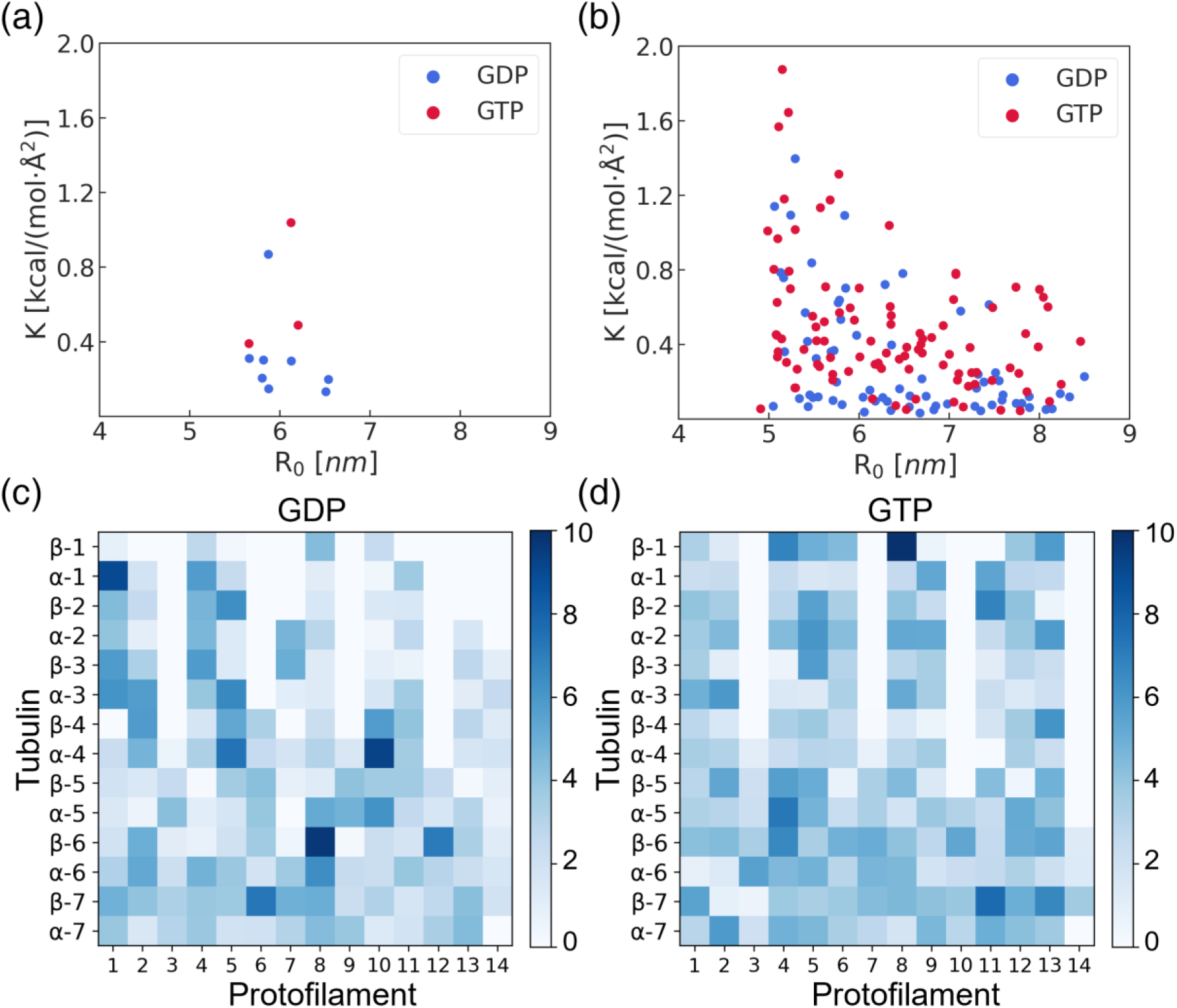
Comparison of the effective interactions between GDP- and GTP-complexed MT tip derived from the HeteroENM analysis. (a) HeteroENM analysis showing the effective interactions of the seam region in both GDP- and GTP-complexed MT states. (b) HeteroENM analysis showing the effective interactions of the non-seam region in both GDP- and GTP-complexed MT states. In both (a) and (b), GDP- and GTP-state effective interactions are shown in *blue* and *red*, respectively. The *x*- and *y*-axes in (a) and (b) represent the length and strength of the effective interaction pairs. (c) The average number of hydrogen bonds formed by a tubulin with its direct right neighboring tubulin in the GDP-complexed MT tip. (d) The average number of hydrogen bonds formed by a tubulin with its direct right neighboring tubulin in the GTP-complexed MT tip. In (c) and (d), the *x*-axis represents the 14 protofilaments, and the *y*-axis shows *α*-or *β*-tubulin of the *i*^th^ heterodimer, arranged from top to the bottom. The color bar indicates the average number of hydrogen bonds shaded from *white* to *blue*.

The number of hydrogen bonds formed by laterally interacting tubulins can reflect the strength of lateral interactions between neighboring PFs. The heatmaps in **Fig. 5*c* and *d*** display the average number of hydrogen bonds formed by a specific tubulin with its direct right neighboring tubulin for the GDP state and GTP state MTs, respectively. The data for this analysis was also collected from the last iteration of the equation-free approach. Variations in the color intensity across the heatmap indicate differences in the number of hydrogen bonds. Generally, the GTP state exhibits darker shades than the GDP state, indicating a greater number of hydrogen bonds on average compared to the lighter shares observed in the GDP state. Each PF of the MT tip in the GDP- or GTP-complexed state may not have the same number of contacts with its neighbors due to the splaying of MT tip. To fairly account for the differences in the number of clusters in the two states, we determined the average number of hydrogen bonds by excluding tubulins that do not have close contacts with their right neighbor. From this analysis, we found the average number of hydrogen bonds per tubulin interacting with its right neighbor to be 3.45 for the GTP state and 2.84 for the GDP state (*p*-value < 0.01 for the one-tailed *T*-test). Interestingly, tubulins located at the bottom of the MT tip do not necessarily form more hydrogen bonds than those at the top. Additionally, the seam region consistently shows fewer hydrogen bonds than the non-seam regions, which is indicated by each rightmost column in **Fig. 5***c* and **5***d*.

## 4. DISCUSSION AND CONCLUDING REMARKS

Conducting sufficiently long simulations to approach the relaxed equilibrium state in extremely large biomolecular systems through all-atom molecular dynamics remains a computational challenge (28,29,65,66), particularly for very large multi-protein complexes such as microtubules. In this work, we employed the multiscale “equation-free” method to accelerate the AA MD relaxation of a 14 PF × 8 heterodimer MT tip lattice. This approach yielded a ∼2-fold computational speedup and demonstrated potential for even greater efficiency under optimal conditions with a loose error threshold. This advance partly addresses a crucial gap in the field, as approaching relaxed structures of this has been a persistent issue (28,29).

Our “equation-free” multiscale approach enabled savings of 0.875 μs all-atom simulation time, which for the two large, many-protein MT system containing ∼21-38 million atoms equate to savings of ∼15 million CPU hours. With an initial 4 μs long AA MD trajectory, we realized near-equilibrium conformations of both GDP- and GTP-complexed MT tips in an effective simulation time of 5.875 μs.

Our analysis of the resulting conformations confirmed differences in the behavior of GDP- and GTP-bound MT tips, particularly in terms of lateral interactions. *In perhaps the most important conclusion from the present work*, here MTs in the GDP state display a tendency to split further into more clusters, while MTs in the GTP state prefer to maintain fewer clusters and exhibit collective behavior, as detailed in the RMSD analysis in **Fig. 3**, **Fig. 4*b***, and the snapshots in the **Supporting Information**. This is an important insight, as lateral interactions are known to play a key role in dynamic instability (12). Additionally, both types of MT tips showed splaying behavior, with GTP-bound MTs exhibiting greater flexibility, which may be advantageous for their biological function such as accommodating various conformations during MT growth (29). A key advantage of the “equation-free” approach is that it is an accelerated sampling technique that maintains all-atom detail and permits us to extract molecular insight that may be lost in fully coarse-grained (CG) models.

The limitations in this study are also crucial to appreciate for a better understanding of the dynamics of MT tips. While our model effectively relaxed the key coarse variables {*r*_*i*,*j*_, θ_*i*,*j*_, φ_*i*,*j*_} on average, it is possible that there are other important slowly-evolving variables parameterizing the slow subspace that are not contained within this set. We note that data-driven CV discovery techniques, particularly those designed to determine slow CVs, present a means to learn CVs from simulation trajectories (49). Furthermore, the high computational cost has limited us to conduct only one independent replica for each of the GDP- and GTP-bound MT tips. As such, we cannot assess how representative each trajectory is of the stochasticity and ensemble of possible dynamical relaxations and terminal relaxed states. Igaev and Grubmüller performed five independent runs for 14 PF ×6 heterodimers long MT tips in GDP- and GTP-bound state (29). Even for the MT tip in the same nucleotide state, the RMSD curves of the five runs can vary differently, indicating that the MT tips are quite flexible and have multiple equilibrium conformations. In our simulations, the MT tips are represented in a simplified manner, with each PF containing only eight tubulin heterodimers, which does not fully capture the complexity of actual MT tips that can consist of hundreds of layers of heterodimers per PF. As Gudimchuk and McIntosh (10) summarized, MT tips can exhibit a large amount of different conformations. Again, the simplifications made here are necessary to make the computational problem tractable within our available computational resources. Additionally, the homogeneity of our model also introduces limitations. In real biological systems, MT tips can exhibit a high degree of flexibility and heterogeneity, such as the ruggedness and variability in the number of tubulin heterodimers per PF among MT tips. This heterogeneity can significantly influence the dynamic instability and mechanical properties of MTs by affecting the interactions of free tubulin heterodimers poised for polymerization. Our current model does not account for this heterogeneity, potentially oversimplifying the dynamic interactions and stability characteristics of MT tips.

Despite these limitations, our current AA MD simulations are designed to accurately model the important molecular interactions at the MT tips, and the currently unprecedented timescale 5.875 μs trajectories have approached the relaxed configurational state and exposed subtle structural differences and the role of lateral interactions in the dynamical relaxations terminal relaxed states of the GDP- and GTP-bound systems. The long AA MD simulation trajectories generated in this work could also potentially serve as a reference trajectory for the development of more sophisticated bottom-up CG models. These future CG models, with a resolution of ∼20 residues per CG site or “bead” (or finer) and encompassing hundreds of layers of tubulins, would enable access to significantly larger system sizes and the possibility of conducing multiple independent CG MD replicas of each system, albeit at lower resolution. We are particularly interested in the potential of CG models to facilitate the study of MT dynamics at a scale closer to biological reality and to incorporate the observed heterogeneity of natural MTs, allowing us to further investigate the MT dynamics and more deeply probe the origin(s) of dynamical instability. The development such models marks the next frontier in our ongoing efforts to simulate and comprehend the complex behavior of MTs in biological systems as well as other many-protein assemblies (67).

## SUPPORTING MATERIAL

Supporting Information can be found online.

## AUTHOR CONTRIBUTIONS

J.W and S.D. contributed equally to this work. G.A.V., A.L.F., and T.C.B. conceptualized and supervised the research. J.W., S.D., and D.B. performed the simulations and wrote the initial draft of the manuscript. All the authors have analyzed the data and contributed to the final version of the manuscript.

## ACKNOWLEDGMENTS

This work was supported by the US Department of Energy, Office of Science, Basic Energy Sciences, under award #DE-SC0023318. We gratefully acknowledge computing time provided by the University of Chicago Research Computing Center (https://rcc.uchicago.edu), the University of Chicago high-performance GPU-based cyberinfrastructure supported by the National Science Foundation under grant no. DMR-1828629, and Frontera at the Texas Advanced Computing Center (TACC) at The University of Texas at Austin (http://www.tacc.utexas.edu) funded by the NSF (OAC-1818253), and the NIH-funded Beagle3 HPC cluster (Award Number S10OD028655).

