## Supporting Information for "Data-Driven Equation-Free Dynamics Applied to Many-Protein Complexes: The Microtubule Tip Relaxation"

### I. The evolution of key MT parameters associated with MT tip splaying

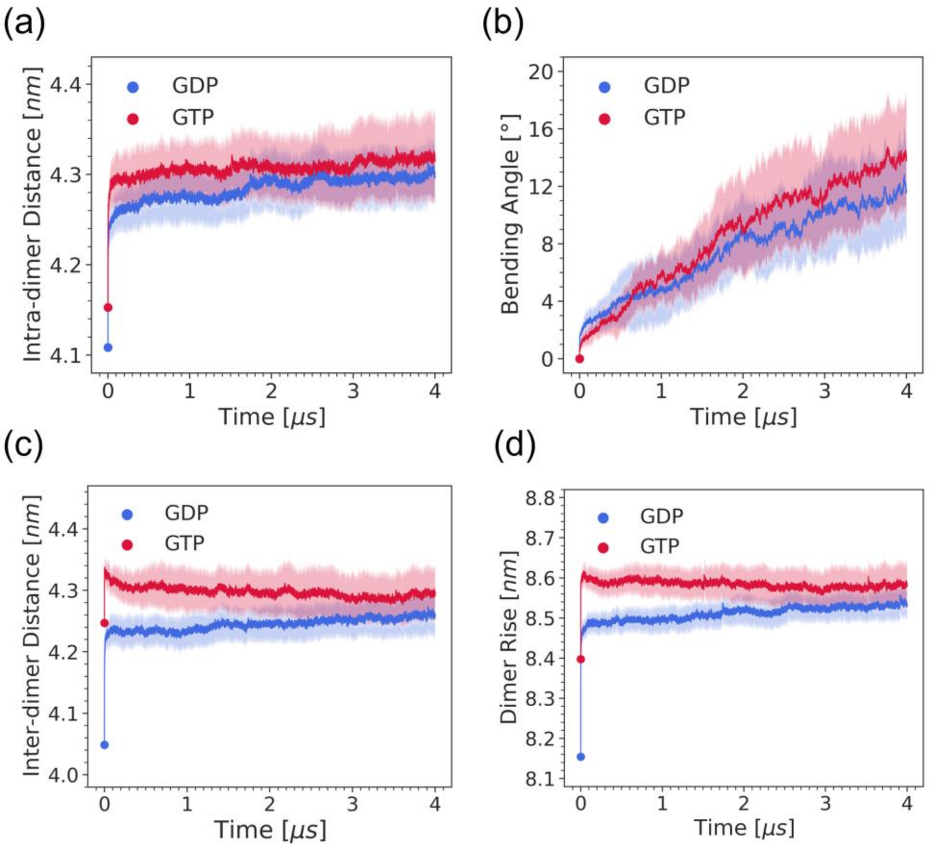


**Figure S1**. **Evolution of key parameters associated with MT Tip splaying process.** (a) Intra-dimer distance, (b) bending angle, (c) inter-dimer distance, and (d) dimer rise as a function of simulation time throughout the initial 4 μs long simulations, respectively, with the GDP-bound MT colored *blue* and GTP-bound MT colored *red*. The solid line indicates the average value over all the 14 PFs and the shade region is the corresponding standard deviation. During the 4 μs all-atom simulation, the bending angle and the intra-dimer distance show an ongoing tendency of increasing, while the inter-dimer distance and dimer rise have a small tendency of either increasing or decreasing with simulation time.

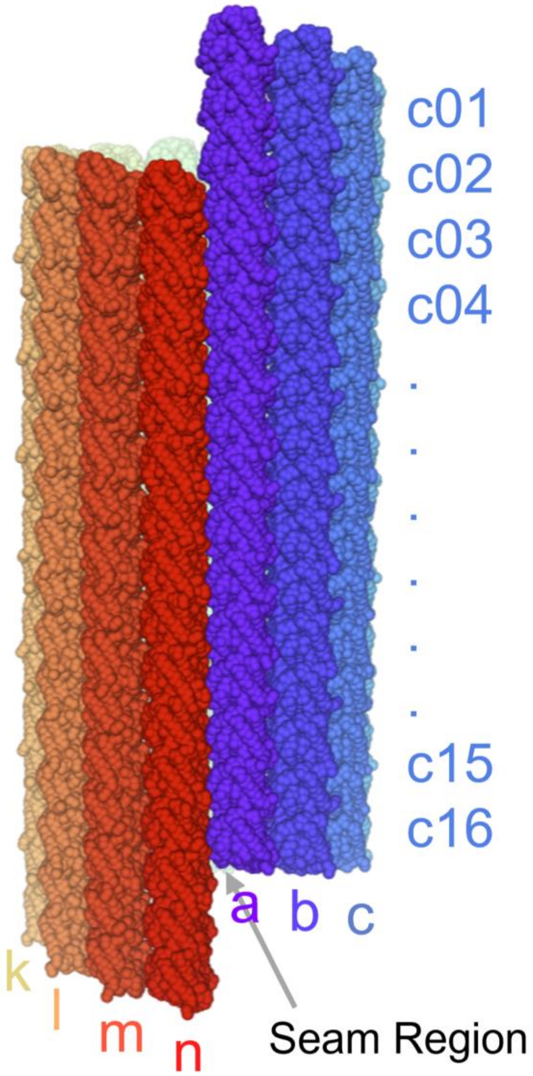


**Figure S2**. **Illustration of the initial straight structure of the MT tip with lattice 14 PF × 8 heterodimer**. Each PF $i \in \{a..n\}$ is distinctly colored from *purple* to *red*. Each tubulin $j \in \{1..16\}$ within a PF *i* is labeled as 01, 02, 03, …, 16, starting from the topmost to the bottommost tubulin in the head-to-tail manner. The $\alpha\beta$-tubulin heterodimers are stacked longitudinally to form protofilaments. The position restraints are applied to the bottommost $\alpha$-tubulins a16-n16 to mimic the MT minus-end. Labeling of PFs and tubulins shown here are employed throughout the main text and the following figures.

**Table S1. Components of the GDP- and GTP-complexed MT tip simulation systems.** "Initial" and "Final" correspond to the simulation system with initial straight cylindrical MT structure and final splaying MT structure, respectively.

| | MT protein atoms | Water molecules | Ion atoms | Total atoms |
|---|---|---|---|---|
| GDP (Initial) | 1,513,680 | 6,309,337 | 41,744 | 20,483,435 |
| GDP (Final) | 1,513,680 | 11,202,205 | 68,982 | 35,189,277 |
| GTP (Initial) | 1,514,240 | 6,475,037 | 42,536 | 20,981,887 |
| GTP (Final) | 1,514,240 | 12,287,759 | 75,006 | 38,452,523 |

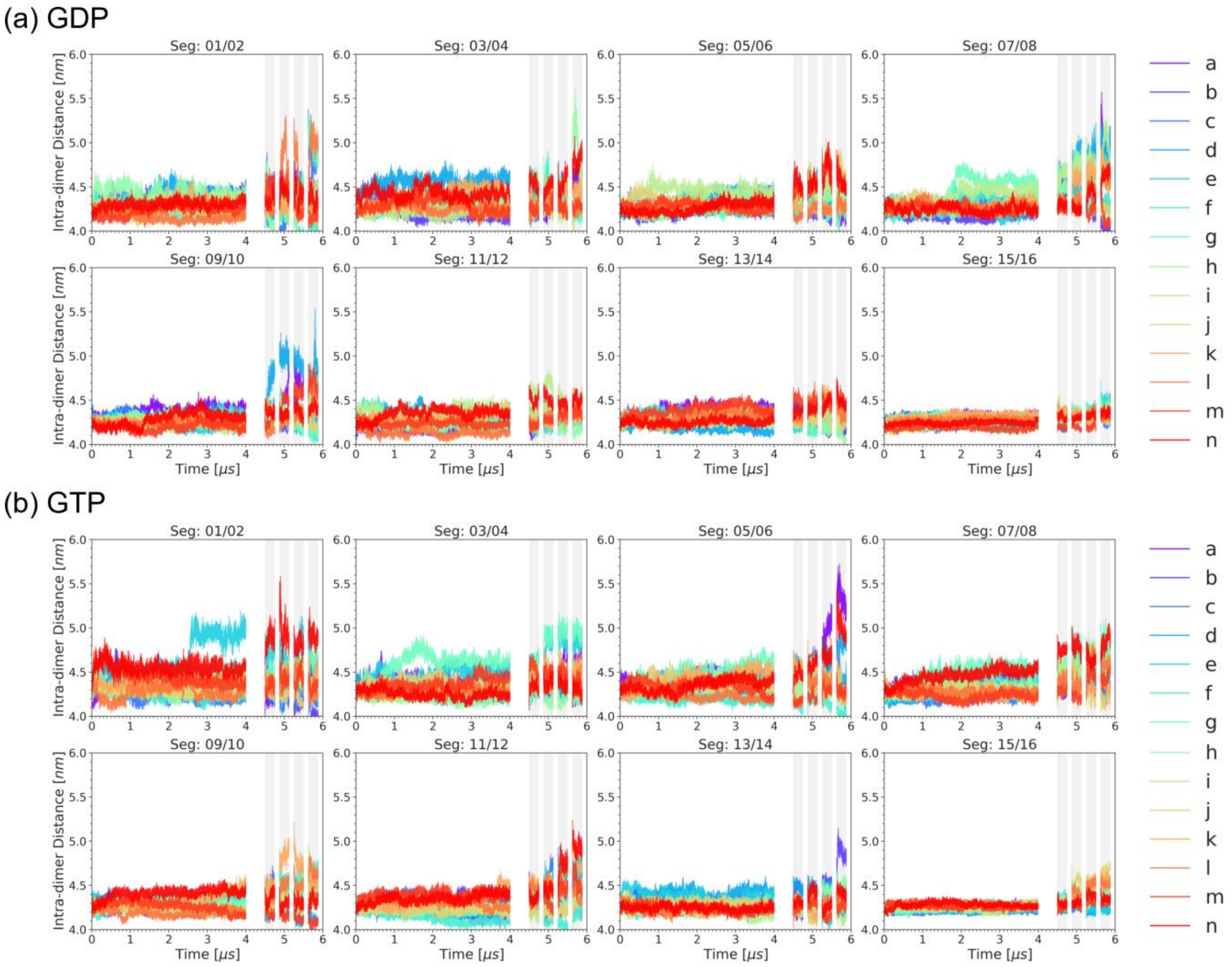


**Figure S3**. **Evolution of intra-dimer distance in the MT tip system.** Intra-dimer distance of (a) GDP-bound state, and (b) GTP-bound state throughout the whole simulation including the samples from the "equation-free" iterations. Each panel represents the intra-dimer distance of a specific layer of MT tip in different PFs. For example, "Seg: 01/02" panel refers to the intradimer distance between tubulin 1 and tubulin 2 in the 14 PFs. The coloring scheme of PFs is consistent with **Fig. S2.** The *gray* regions denote the four equation-free iterations from 1 to 4, respectively.

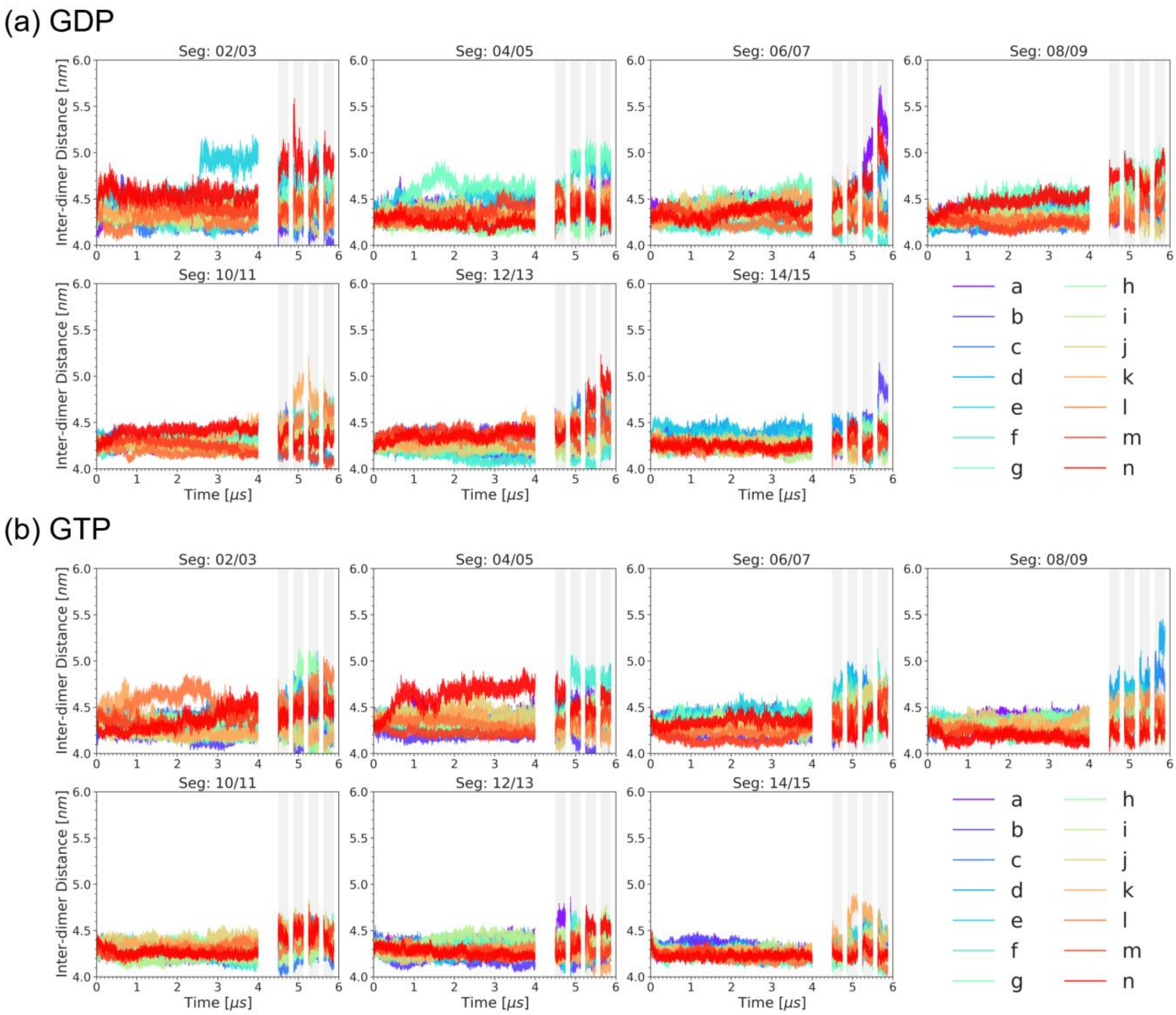


**Figure S4**. **Evolution of inter-dimer distance in the MT tip system.** Inter-dimer distances of (a) GDP-bound state, and (b) GTP-bound state throughout the whole simulation including the samples from the "equation-free" iterations. Each panel represents the inter-dimer distance of a specific layer of MT tip in different PFs. For example, "Seg: 02/03" panel refers to the inter-dimer distance between tubulin 2 and tubulin 3 in the 14 PFs. The coloring scheme of PFs is consistent with **Fig. S2.** The *gray* regions denote the four equation-free iterations from 1 to 4, respectively.

### II. Convergence of bending and twisting motions with "equation-free" iterations

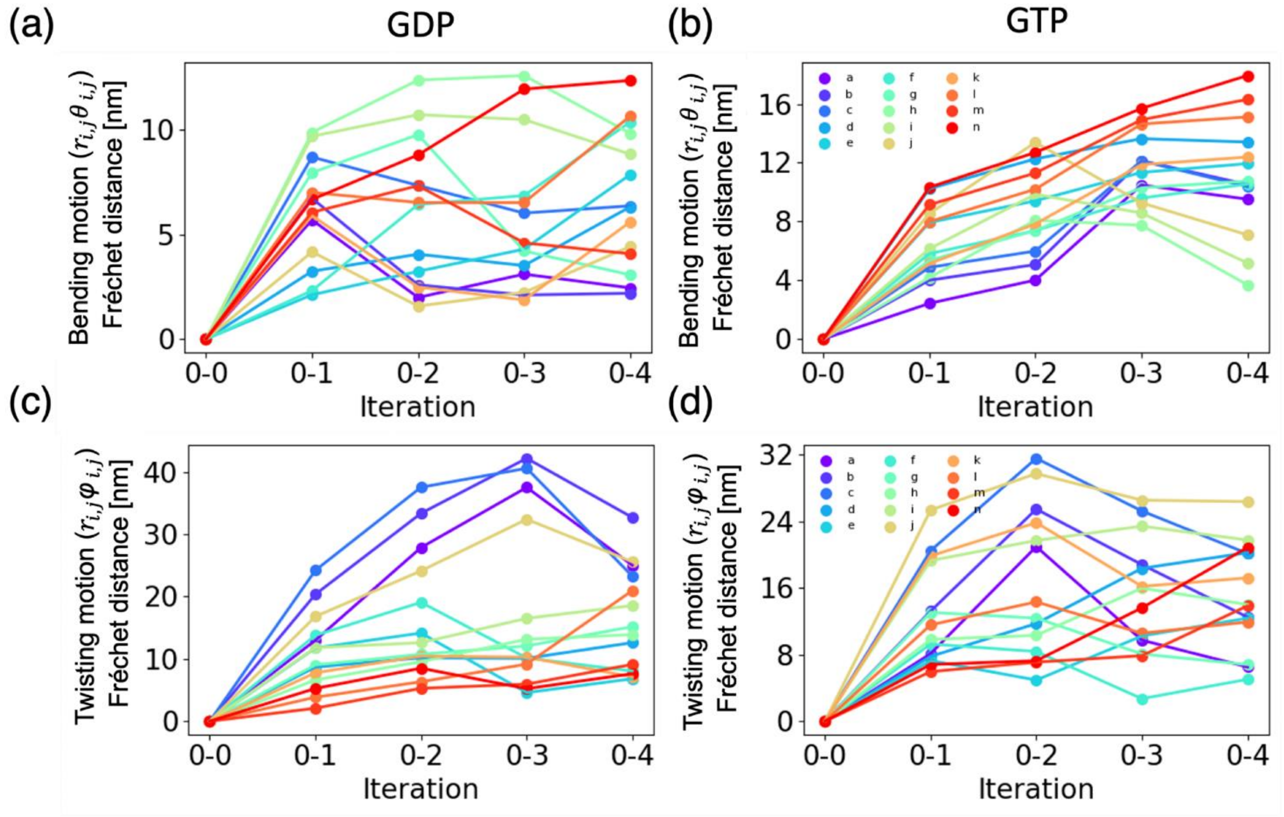


**Figure S5**. **Variation of Fréchet distance between bending and twisting curves in an iteration of the "equation-free" method and that observed in the initial 4 μs trajectory.** (a) Changes in the Fréchet distance between the 16 time-averaged $r_{i,j}$-$\theta_{i,j}$ coordinates in each of the 14 PFs in each of the iteration with respect to their corresponding coordinates in the initial 4 μs trajectory for GDP-complexed MT tip system, (b) Similar to (a) but for the GTP-complexed MT tip system. (c) Changes in the Fréchet distance between the 16 time-averaged $r_{i,j}$-$\varphi_{i,j}$ coordinates in each of the 14 PFs in each of the iteration with respect to their corresponding coordinates in the initial 4 μs trajectory for GDP-complexed MT tip system, (d) Similar to (c) but for the GTP-complexed MT tip system. In (a)-(d), the Fréchet distances corresponding to each PF are plotted in different colors with the coloring scheme consistent with **Fig. S2**.

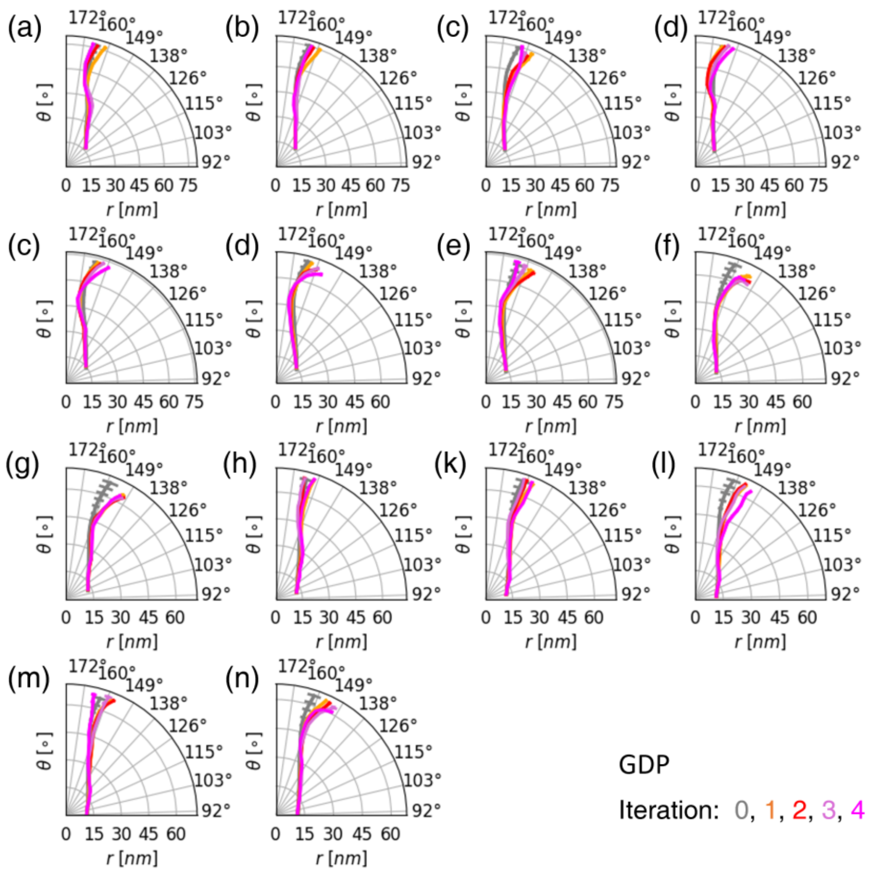


**Figure S6**. **Bending profiles of the PFs in the GDP-complexed MT tip system observed in the initial 4 μs trajectory and the four "equation-free" method iterations.** (a)-(n) Plots of the 16 time-averaged ($r_{i,j}$,$\theta_{i,j}$) coordinates corresponding to the 16 tubulins $j$, where each panel corresponds to one of the PF $i \in \{a..n\}$. In (a)-(n), the time averaged profile observed in the initial 4 μs trajectory is shown using grey solid line. The tangential error bar indicates the standard deviation of $\theta_{i,j}$ observed during the simulation. Time averaged bending profiles along with the standard deviations observed in the four "equation-free" iterations are colored in orange, red, purple, and magenta, respectively.

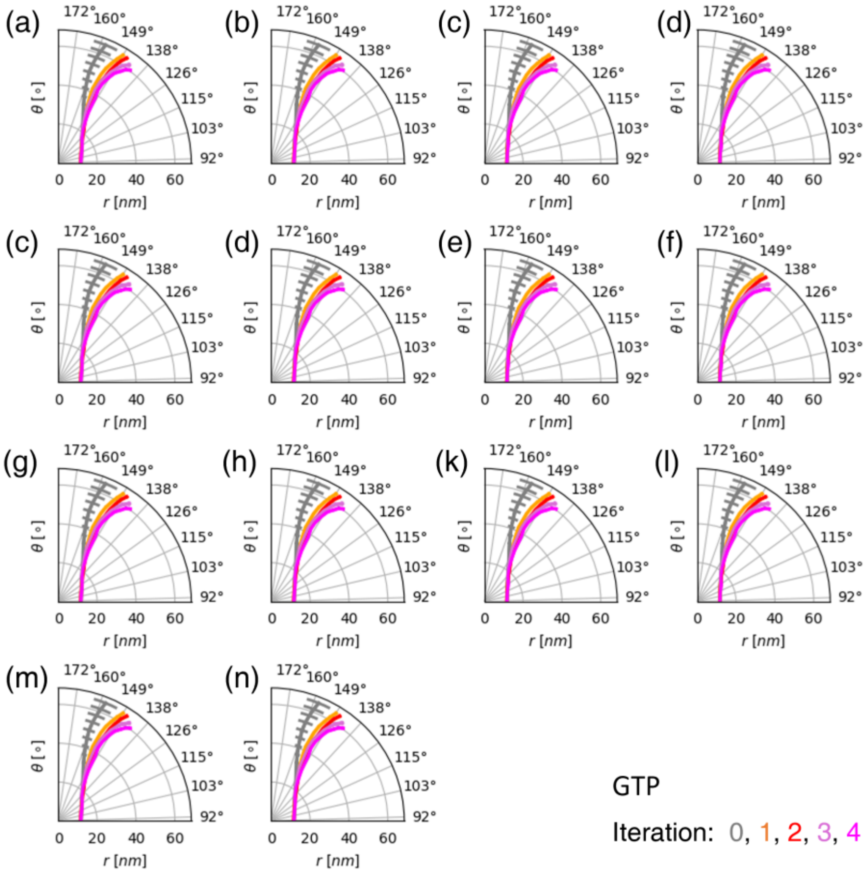


**Figure S7**. **Bending profiles of the PFs in the GTP-complexed MT tip system observed in the initial 4 μs trajectory and the four "equation-free" method iterations.** (a)-(n) Plots of the 16 time-averaged $(r_{i,j},\theta_{i,j})$ coordinates corresponding to the 16 tubulins $j$, where each panel corresponds to one of the PF $i \in \{a..n\}$. In (a)-(n), the time averaged profile observed in the initial 4 μs trajectory is shown using grey solid line. The tangential error bar indicates the standard deviation of $\theta_{i,j}$ observed during the simulation. Time averaged bending profiles along with the standard deviations observed in the four "equation-free" iterations are colored in orange, red, purple, and magenta, respectively.

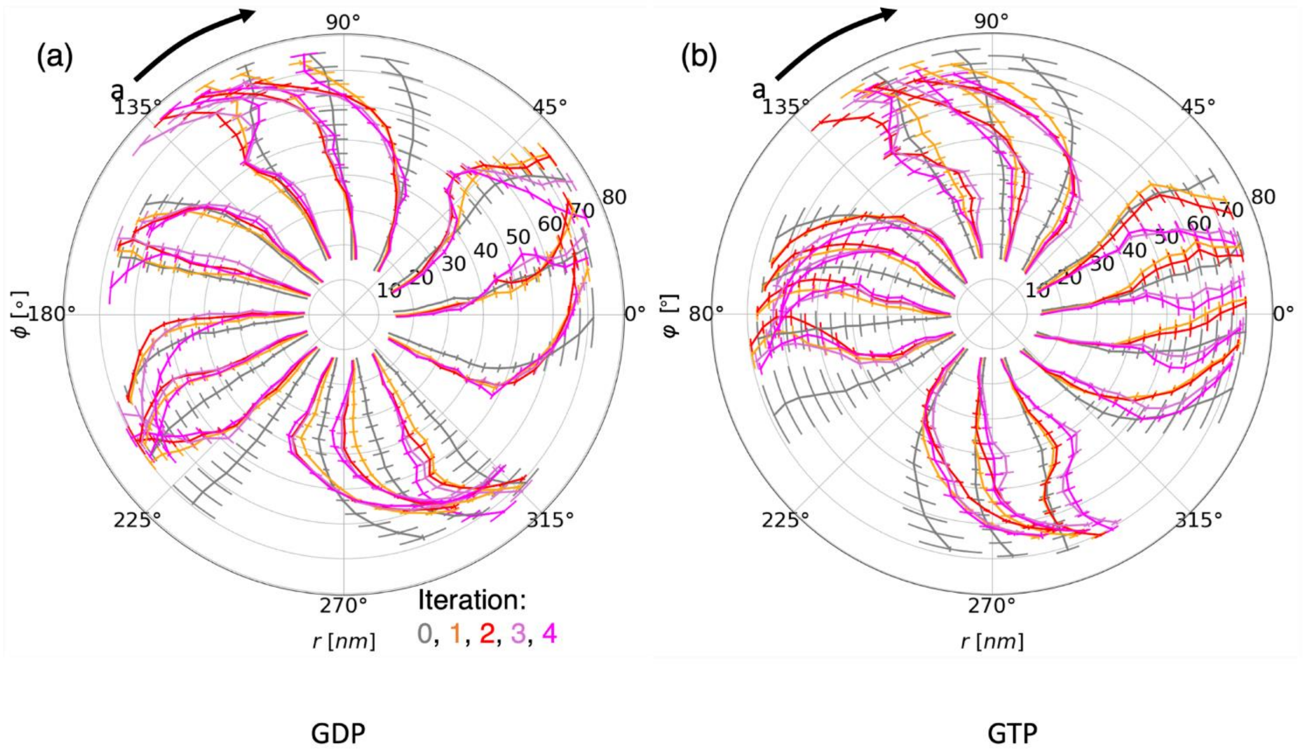


**Figure S8**. **Twisting profiles of the PFs in the GDP- and GTP- complexed MT tip system observed in the initial 4 μs trajectory and the four "equation-free" method iterations.** (a) Plots of the 16 time-averaged ($r_{i,j}$,$\varphi_{i,,j}$) coordinates corresponding to the 16 tubulins $j$ in GDP-complexed MT tip system. Each solid line corresponds to one of the PF $i \in \{a..n\}$. The time averaged profile observed in the initial 4 μs trajectory is shown using grey solid line. The tangential error bar indicates the standard deviation of $\varphi_{i,j}$ observed during the simulation. Time averaged bending profiles along with the standard deviations observed in the four "equation-free" iterations are colored in orange, red, purple, and magenta, respectively. (b) Similar to (a) but for the GTP-complexed MT tip system.

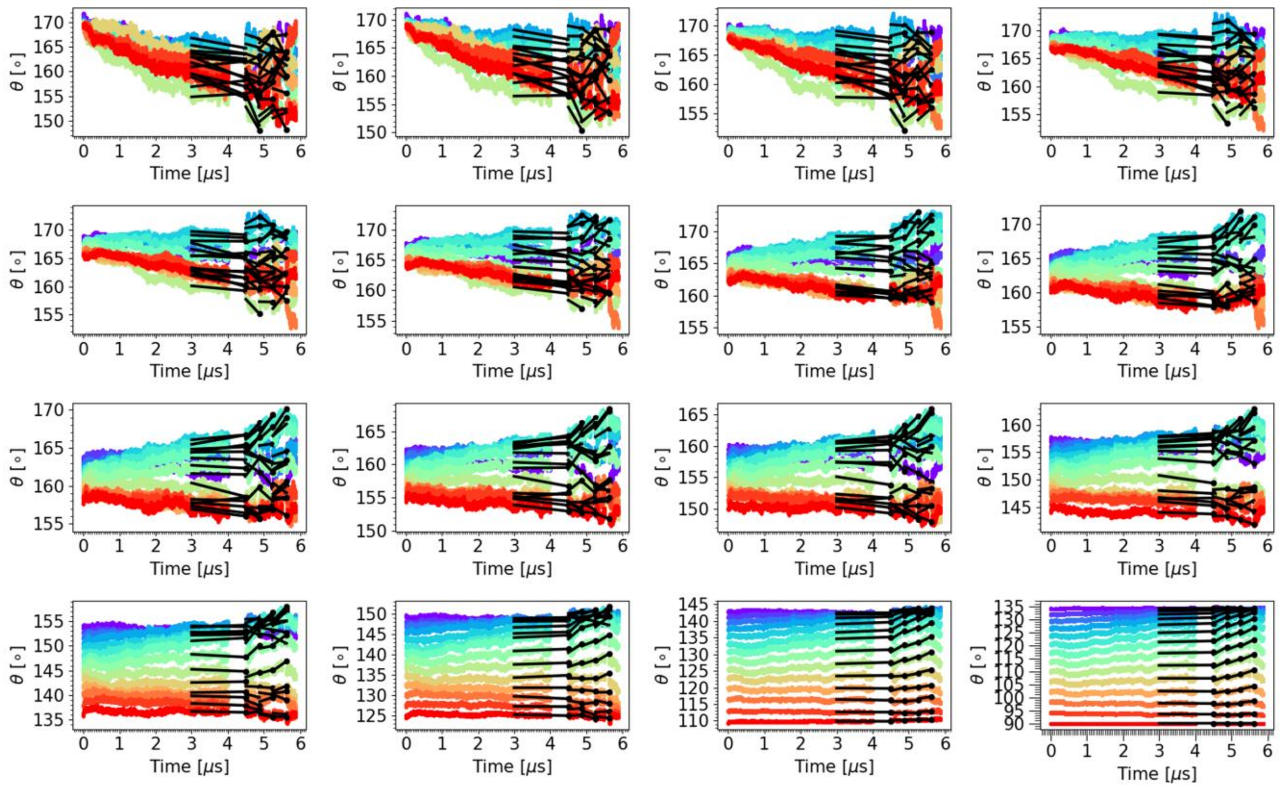


**Figure S9**. **Evolution of $\theta_{i,j}$ CVs in the GDP-complexed MT tip system during the "equation-free" method.** Each panel represents the changes in $\theta_{i,j}$ of different PFs $i$ in a tubulin $j$. The panels from top left to bottom right correspond to the 16 tubulins arranged from top to the bottom layer. For example, the first panel represents the first (uppermost) layer, which consists of $\beta$-tubulins. The last panel represents the bottommost layer composed of $\alpha$-tubulins. Different colors represent different values of $i$ corresponding to different PFs $i \in \{a..n\}$. The coloring code is consistent with **Fig. S2**. Solid black lines in each plot represent the linear models applied during the projection step of the "equation-free" method. Solid black circle markers indicate the projected or predicted value of $\theta_{i,j}$ in each iteration that was used during the lifting operation for initiating a new short all-atom MD simulation.

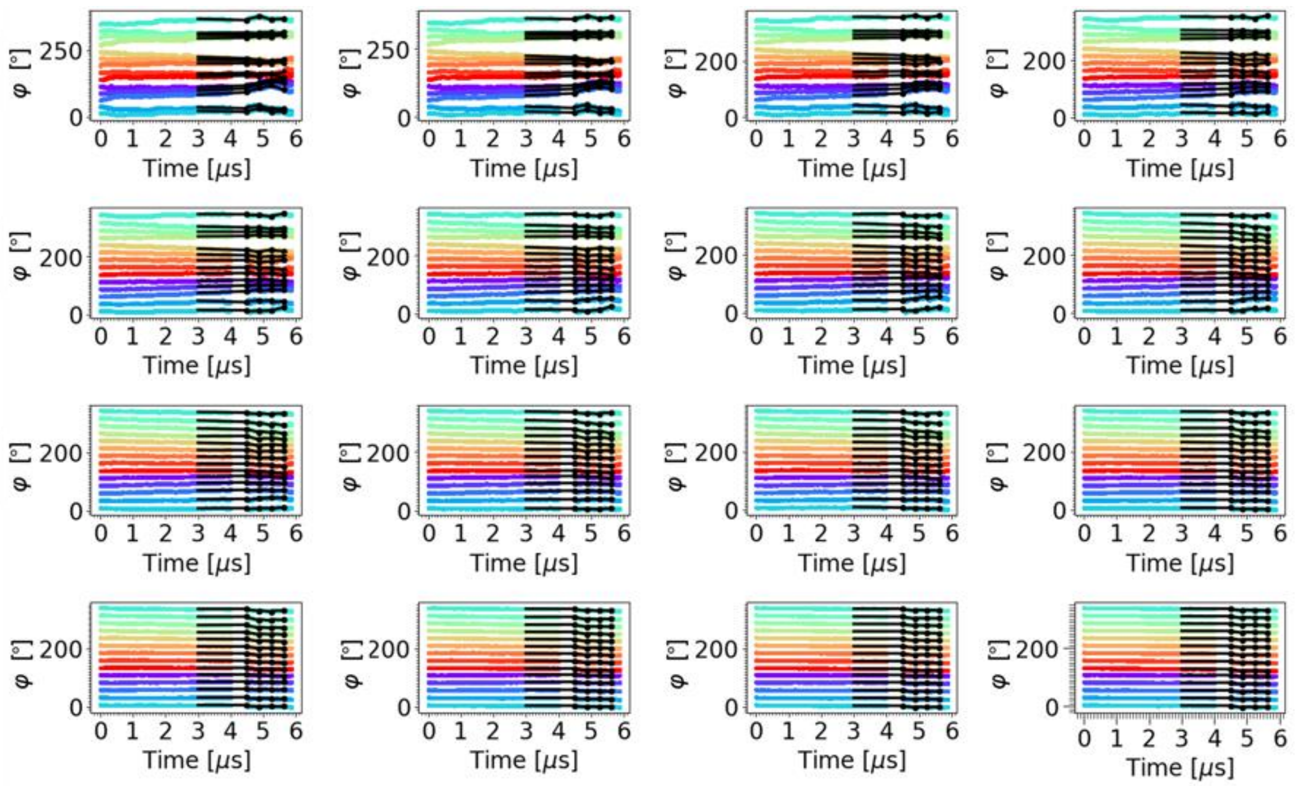


**Figure S10**. **Evolution of $\varphi_{i,j}$ CVs in the GDP-complexed MT tip system during the "equation-free" method.** Each panel sequentially represents the changes in $\varphi_{i,j}$ of different PFs $i$ in a tubulin $j$. The panels from top left to bottom right correspond to the 16 tubulins arranged from top to the bottom layer. For example, the first panel represents the first (uppermost) layer, which consists of $\beta$-tubulins. The last panel represents the bottommost layer composed of $\alpha$-tubulins. Different colors represent different values of $i$ corresponding to different PFs $i \in \{a..n\}$. The coloring code is consistent with **Fig. S2**. Solid black lines in each plot represent the linear models applied during the projection step of the "equation-free" method. Solid black circle markers indicate the projected or predicted value of $\varphi_{i,j}$ in each iteration that was used during the lifting operation for initiating a new short all-atom MD simulation.

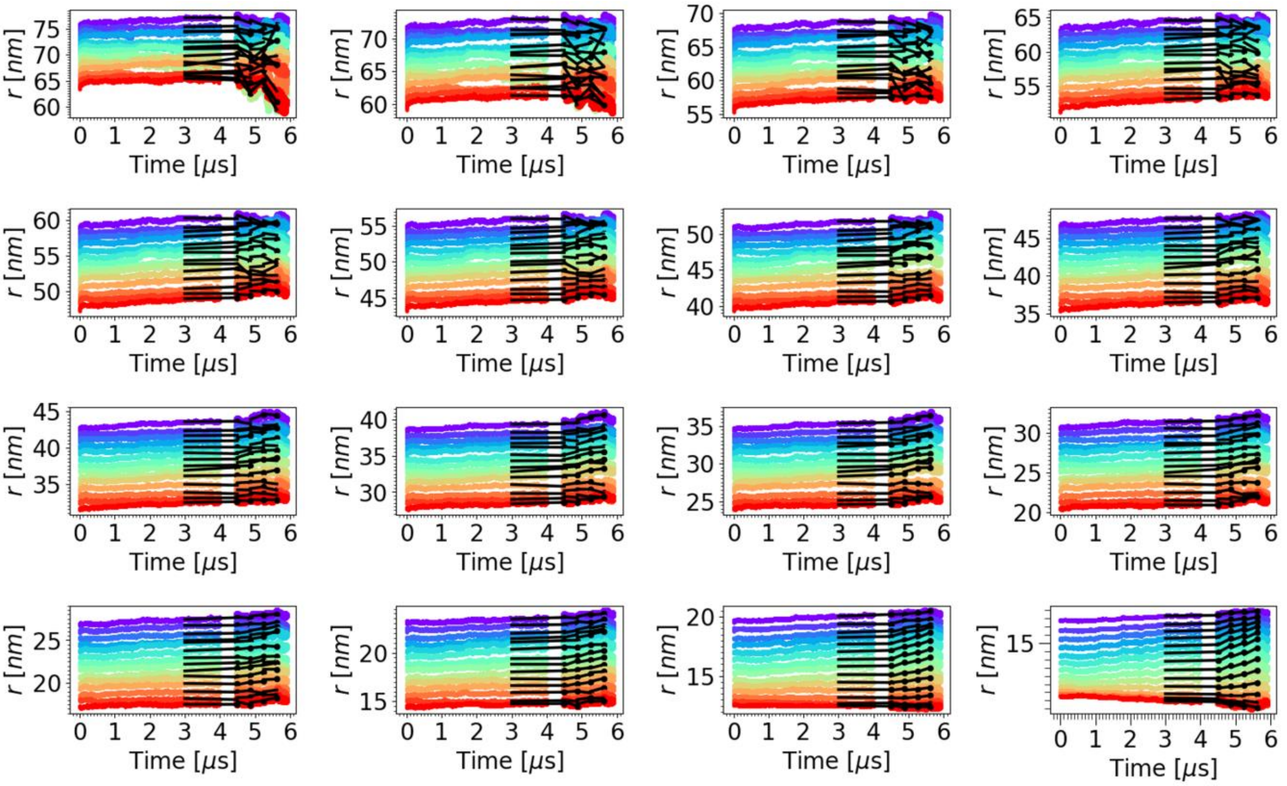

**Figure S11**. **Evolution of r$_{i,j}$ CVs in the GDP-complexed MT tip system during the "equation-free" method.** Each panel sequentially represents the changes in r$_{i,j}$ of different PFs $i$ in a tubulin $j$. The panels from top left to bottom right correspond to the 16 tubulins arranged from top to the bottom layer. For example, the first panel represents the first (uppermost) layer, which consists of $\beta$-tubulins. The last panel represents the bottommost layer composed of $\alpha$-tubulins. Different colors represent different values of $i$ corresponding to different PFs $i \in \{a..n\}$. The coloring code is consistent with **Fig. S2**. Solid black lines in each plot represent the linear models applied during the projection step of the "equation-free" method. Solid black circle markers indicate the projected or predicted value of r$_{i,j}$ in each iteration that was used during the lifting operation for initiating a new short all-atom MD simulation.

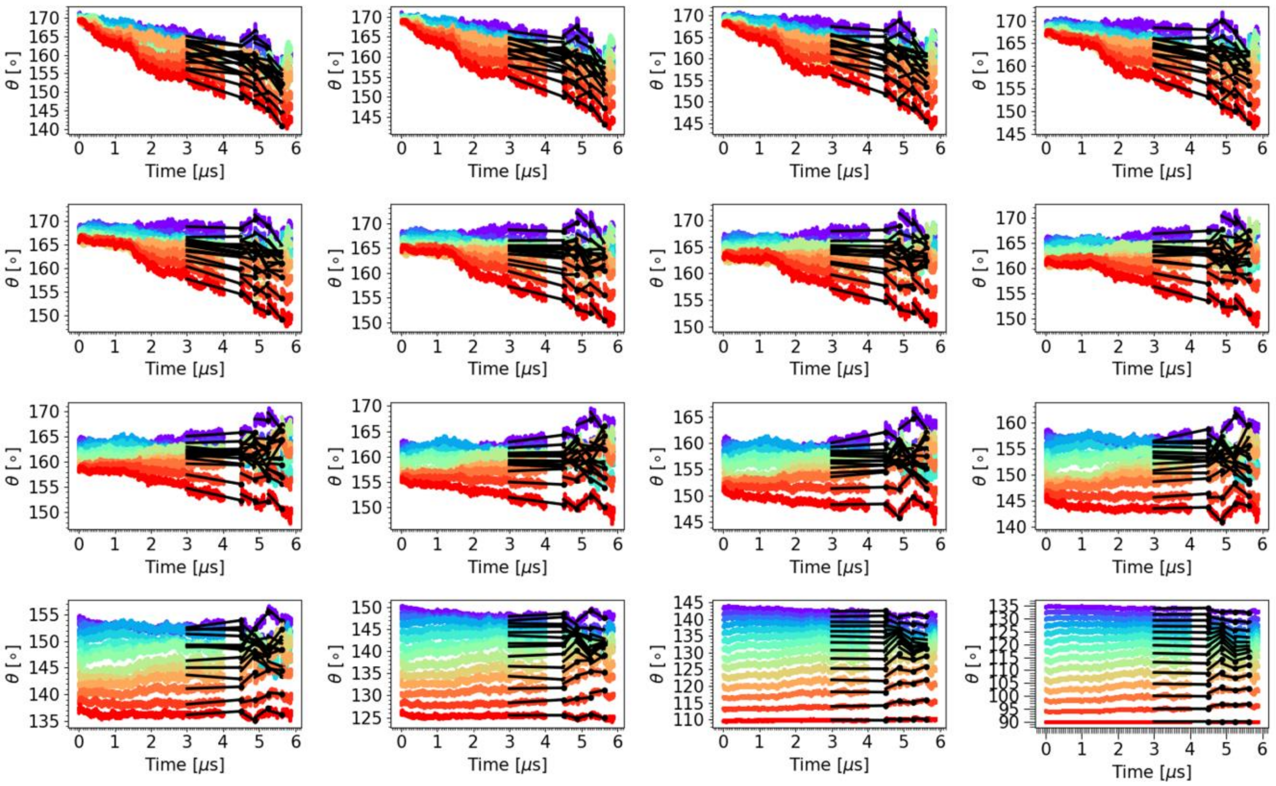


**Figure S12**. **Evolution of $\theta_{i,j}$ CVs in the GTP-complexed MT tip system during the "equation-free" method.** Each panel sequentially represents the changes in $\theta_{i,j}$ of different PFs $i$ in a tubulin $j$. The panels from top left to bottom right correspond to the 16 tubulins arranged from top to the bottom layer. For example, the first panel represents the first (uppermost) layer, which consists of $\beta$-tubulins. The last panel represents the bottommost layer composed of $\alpha$-tubulins. Different colors represent different values of $i$ corresponding to different PFs $i \in \{a..n\}$. The coloring code is consistent with **Fig. S2**. Solid black lines in each plot represent the linear models applied during the projection step of the "equation-free" method. Solid black circle markers indicate the projected or predicted value of $\theta_{i,j}$ in each iteration that was used during the lifting operation for initiating a new short all-atom MD simulation.

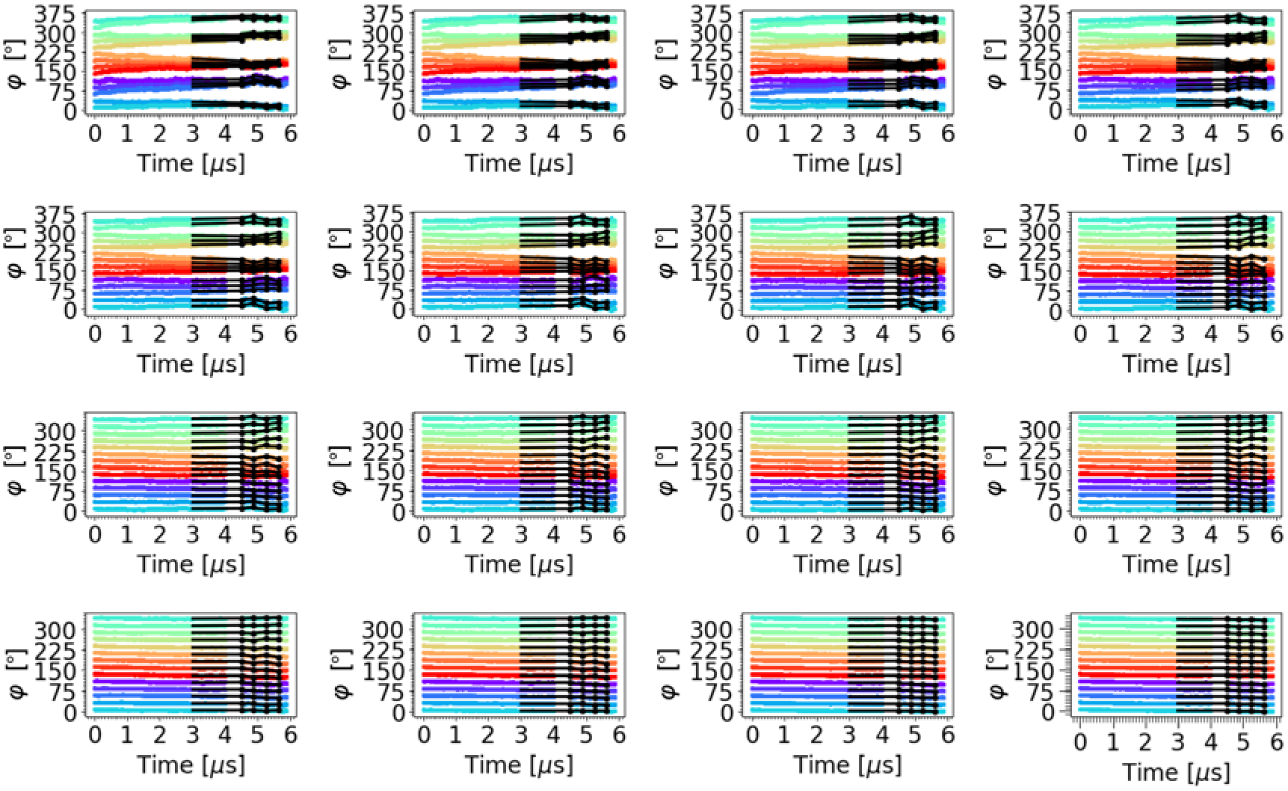


**Figure S13**. **Evolution of $\varphi_{i,j}$ CVs in the GTP-complexed MT tip system during the "equation-free" method.** Each panel sequentially represents the changes in $\varphi_{i,j}$ of different PFs $i$ in a tubulin $j$. The panels from top left to bottom right correspond to the 16 tubulins arranged from top to the bottom layer. For example, the first panel represents the first (uppermost) layer, which consists of $\beta$-tubulins. The last panel represents the bottommost layer composed of $\alpha$-tubulins. Different colors represent different values of $i$ corresponding to different PFs $i \in \{a..n\}$. The coloring code is consistent with **Fig. S2**. Solid black lines in each plot represent the linear models applied during the projection step of the "equation-free" method. Solid black circle markers indicate the projected or predicted value of $\varphi_{i,j}$ in each iteration that was used during the lifting operation for initiating a new short all-atom MD simulation.

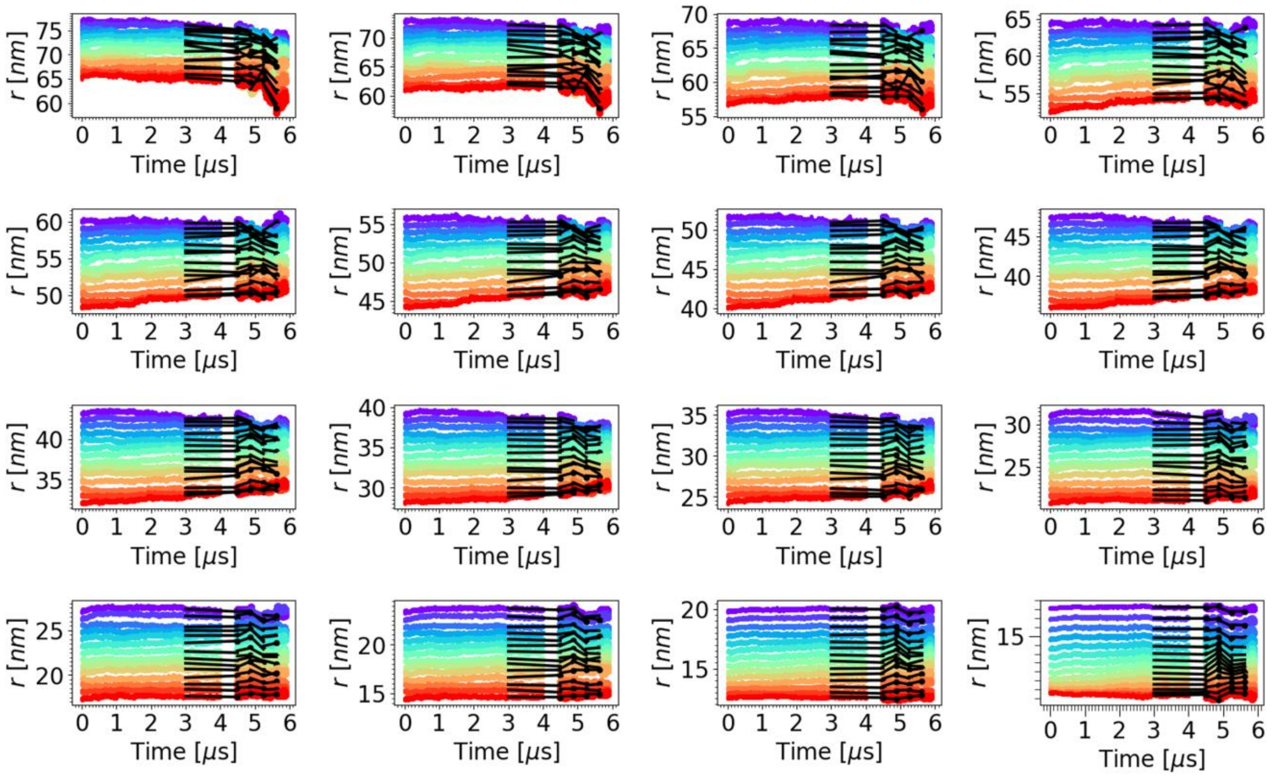


**Figure S14**. **Evolution of r$_{i,j}$ CVs in the GTP-complexed MT tip system during the "equation-free" method.** Each panel sequentially represents the changes in r$_{i,j}$ of different PFs $i$ in a tubulin $j$. The panels from top left to bottom right correspond to the 16 tubulins arranged from top to the bottom layer. For example, the first panel represents the first (uppermost) layer, which consists of $\beta$-tubulins. The last panel represents the bottommost layer composed of $\alpha$-tubulins. Different colors represent different values of $i$ corresponding to different PFs $i \in \{a..n\}$. The coloring code is consistent with **Fig. S2**. Solid black lines in each plot represent the linear models applied during the projection step of the "equation-free" method. Solid black circle markers indicate the projected or predicted value of r$_{i,j}$ in each iteration that was used during the lifting operation for initiating a new short all-atom MD simulation.

### III. Snapshots illustrating the splaying process of MT tip system

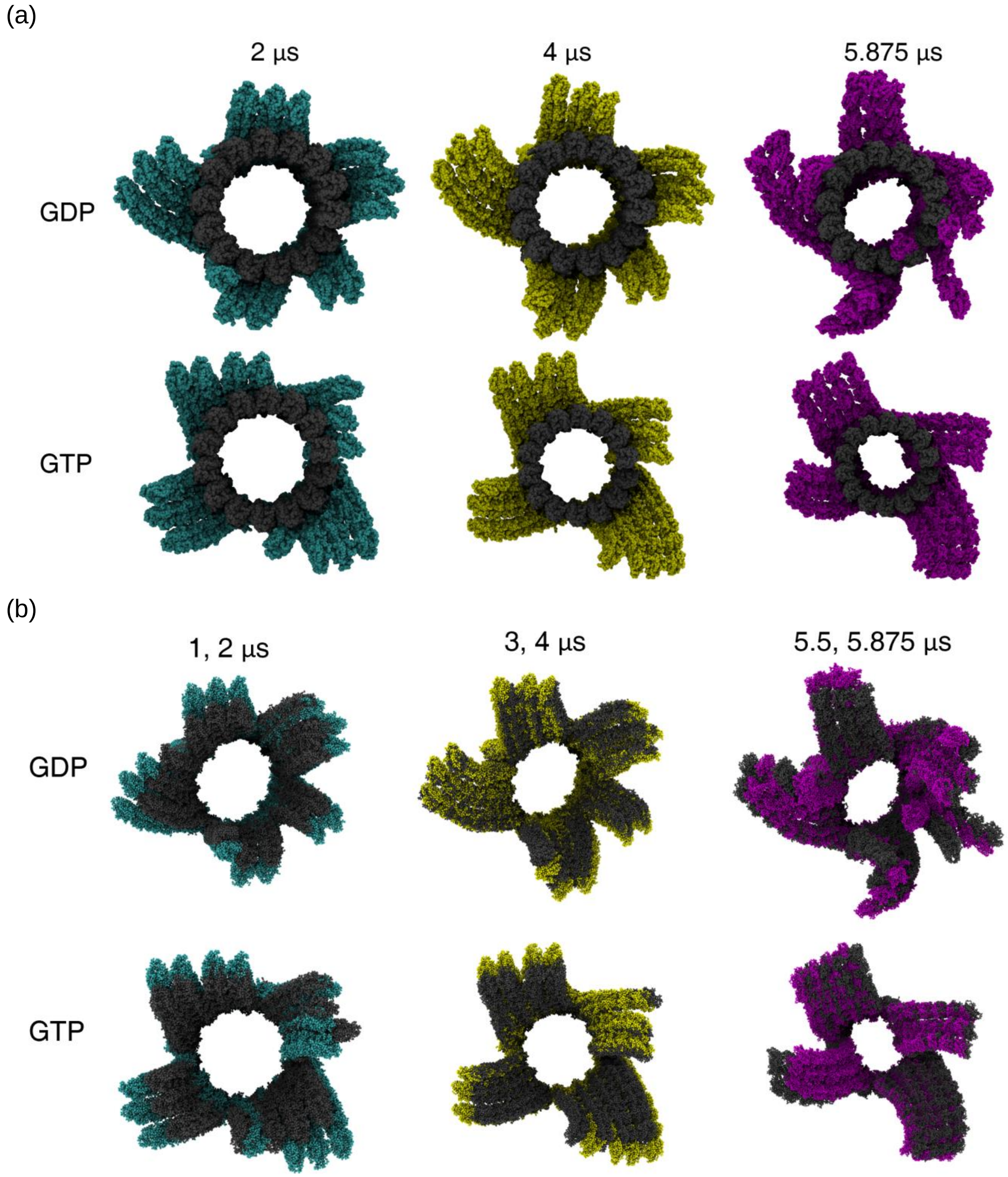


**Figure S15**. **Visual analysis of the splaying process of GDP- and GTP- complexed MT tip system**. (a) Snapshots (top view) for MTs in the GDP- and GTP-bound states at 2 μs, 4 μs, and 5.875 μs, with the *grey* structures representing the straight lattice as a reference. (b) Snapshots (top view) for MTs in GDP and GTP states at 2 μs, 4 μs, and 5.875 μs, with the *grey* structures representing the structures at 1 μs, 3 μs, and 5.5 μs for reference, respectively.

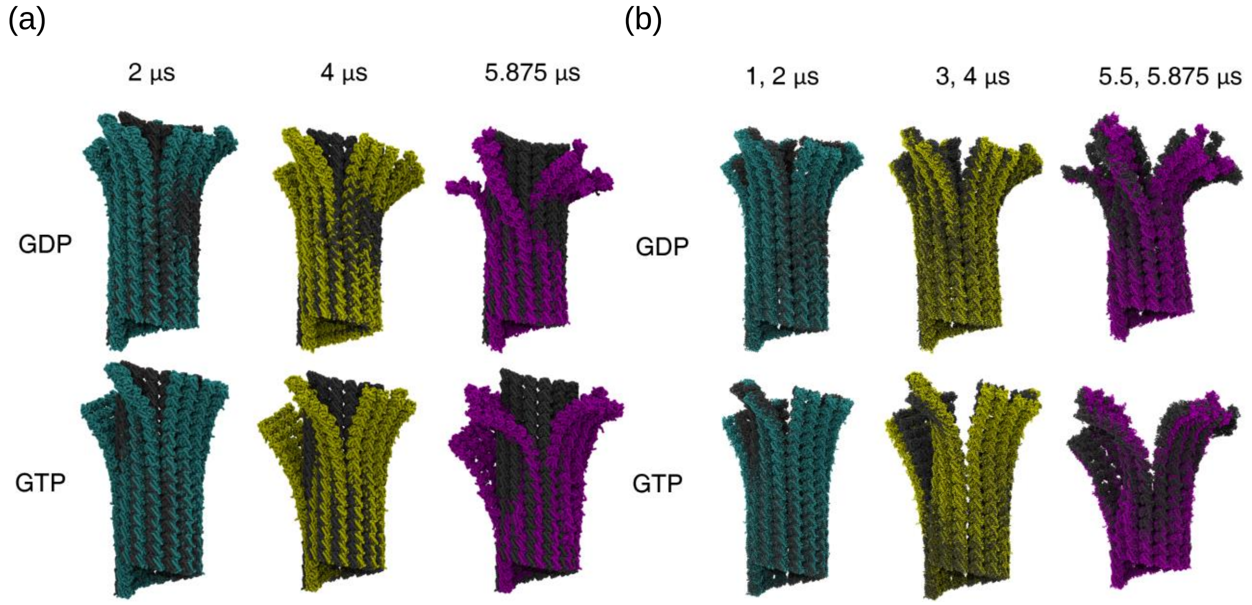


**Figure S16**. **Visual analysis of the splaying process of GDP- and GTP- complexed MT tip system**. (a) Snapshots (side view) for MTs in the GDP- and GTP-bound states at 2 μs, 4 μs, and 5.875 μs, with the *grey* structures representing the straight lattice as a reference. (b) Snapshots (side view) for MTs in GDP and GTP states at 2 μs, 4 μs, and 5.875 μs, with the *grey* structures representing the structures at 1 μs, 3 μs, and 5.5 μs for reference, respectively.

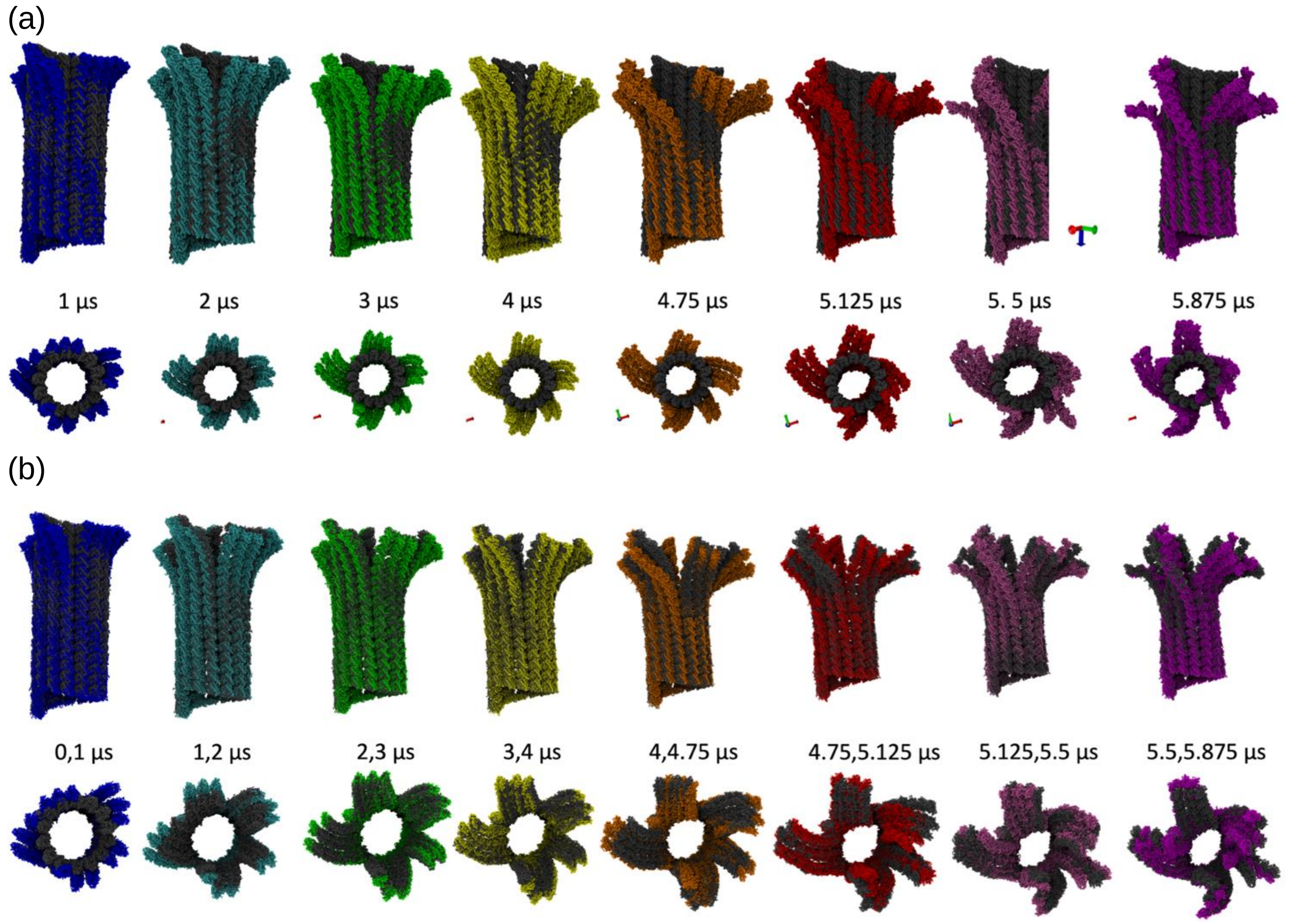


**Figure S17**. **Visual analysis of the splaying process of GDP-complexed MT tip system**. (a) Snapshots (side view) for MTs in the GDP-bound state at 1, 2, 3, 4, 4.75, 5.125, 5.5, 5.875 μs, with the *grey* structures representing the straight lattice as a reference. (b) Snapshots (side view) for MTs in the GDP-bound state, with the *grey* structures representing the configuration at the previous stage for reference. For example, "3, 4 μs" means the colored structure is the conformation at 4 μs and the *grey* structure is the conformation at 3 μs.

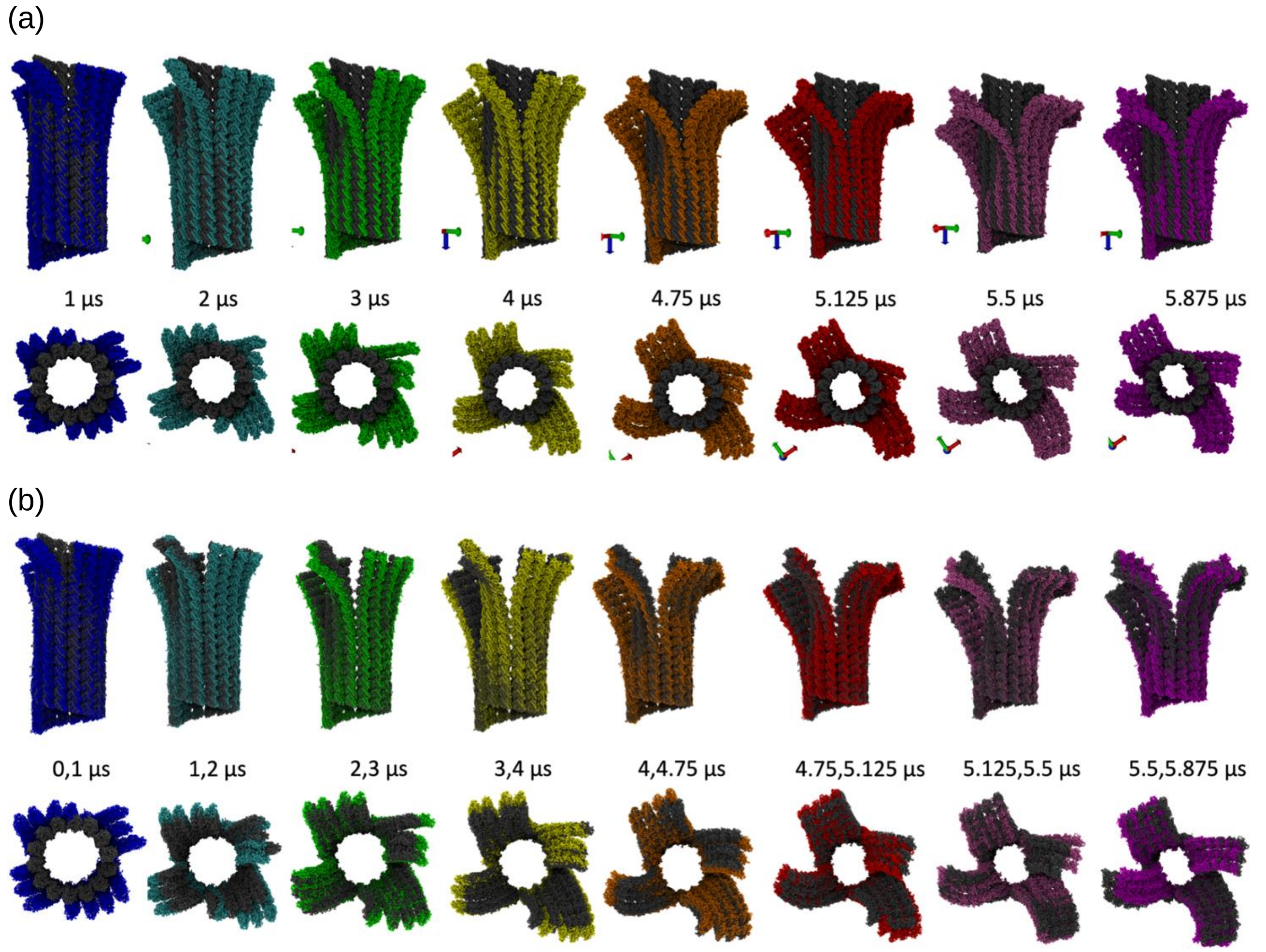


**Figure S18**. **Visual analysis of the splaying process of GTP- complexed MT tip system**. (a) Snapshots (side view) for MTs in the GTP-bound state at 1, 2, 3, 4, 4.75, 5.125, 5.5, 5.875 μs, with the *grey* structures representing the straight lattice as a reference. (b) Snapshots (side view) for MTs in the GTP-bound state, with the *grey* structures representing the configuration at the previous stage for reference. For example, "3, 4 μs" means the colored structure is the conformation at 4 μs and the *grey* structure is the conformation at 3 μs.

### V. Retrospective analysis for calibrating equation-free training time *T* and jump stride *τ*

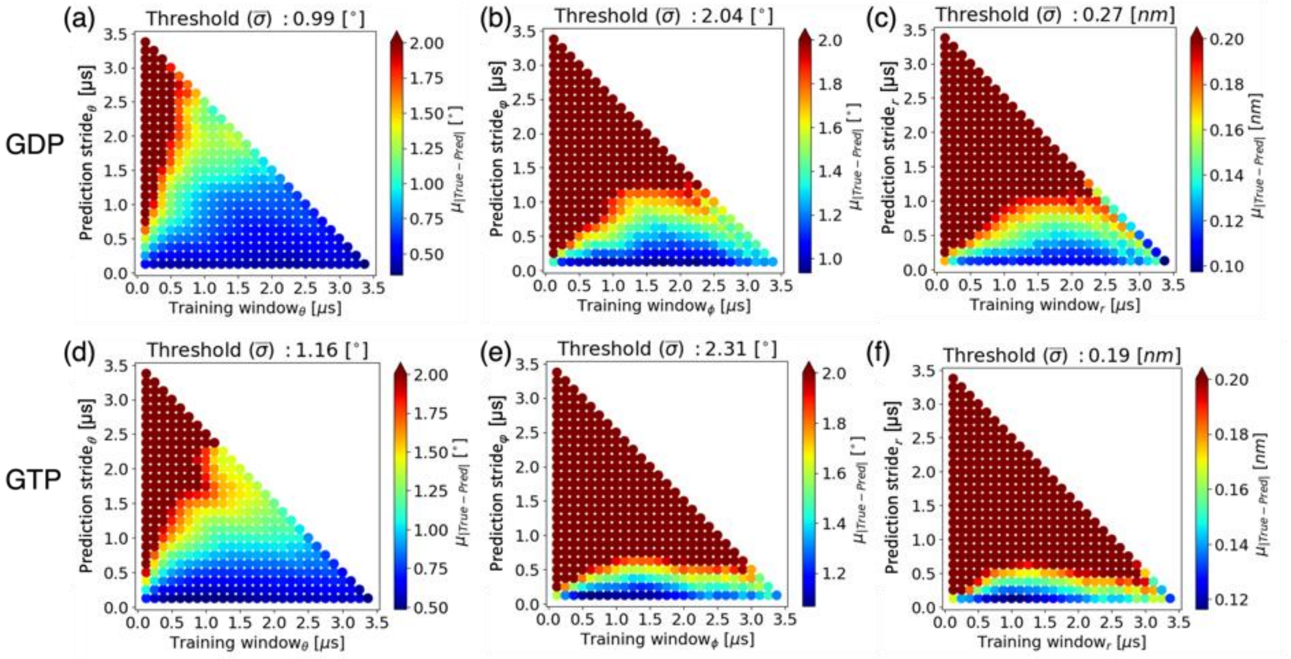


**Figure S19**. **Selection of an appropriate training trajectory window time *T* and coarse-projective jump stride $\tau$ during the "equation-free" projection step.** In each plot, the grid point shows the mean absolute error for a given training window *T* and prediction stride $\tau$, averaged over all 14 × 16 tubulins for the CVs (a) $\theta_{i,j}$, (b) $\varphi_{i,j}$, and (c) $r_{i,j}$ in GDP-complexed MT tip system. Similar profiles for GTP-complexed MT tip system are shown in (d)-(f). Each grid point in (a)-(f) is computed by first training linear models over different trajectory window times *T* using the initial 4 μs trajectory. Then, the absolute error between the predictions using the linear models and the true values observed in the initial 4 μs trajectory are calculated at different jump stride times $\tau$. The mean absolute errors over all given trajectory window times *T* and jump stride times $\tau$ are then calculated from the absolute errors. Average of the mean absolute errors over all 14 × 16 tubulins are then calculated. These averages represent the color of each grid point in (a)-(f). In each plot, the x- and y-axis show the trajectory window times *T* and jump stride times $\tau$. Above each panel, the threshold for the maximum error tolerance is shown. We set the threshold of each CV to the standard deviation of the CVs averaged over all 14 × 16 tubulins ($\bar{\sigma}$) in the initial 4 μs trajectory. Although larger training window and smaller prediction stride can give more accurate predictions (*blue*), the computational speedup is also reduced. For this reason, we selected a prediction stride $\tau$ and training window time *T* that significantly boosts the computational performance while keeping the potential prediction error within the threshold and judiciously utilizing the computational resources required for the entire "equation-free" iterative process.
